# Global Regulatory Functions of the Human CCR4–NOT Complex in mRNA Deadenylation and Decay

**DOI:** 10.64898/2026.09.30.755734

**Authors:** Katherine M. McKenney, Mika Nevo, Robert P. Connacher, Khadija Rebbani, Zijun Luo, Elise B. Dunshee, Marc R. Fabian, Eugene Valkov, Justin Brumbaugh, Aaron C. Goldstrohm

## Abstract

Shortening of the poly(A) tail initiates mRNA decay, and the human CCR4–NOT is the principal cytoplasmic deadenylase. Although its subunits have been defined, their functions have not been comprehensively evaluated in cells. We depleted each CCR4–NOT subunit and measured transcriptome-wide effects. CCR4–NOT regulated transcripts from thousands of genes, many of which associated with the complex in cells. Distinct groups of transcripts responded according to the modular organization of the complex, while subsets exhibited unique responses to specific subunits. Poly(A) tail analysis revealed that the CNOT7 and CNOT8 deadenylases are the predominant regulators of poly(A) tail length in these cells. CCR4–NOT targets were enriched in regulatory pathways including cell proliferation, apoptosis, stress responses, and cancer-related processes. Bioinformatic analysis identified sequence determinants and candidate recruitment factors associated with CCR4–NOT regulation. The results define the breadth and specificity of CCR4–NOT-mediated mRNA decay and provide a resource for dissecting regulatory mechanisms and biological roles.

**Graphical Abstract:** CCR4–NOT functions as a modular mRNA regulatory platform that links deadenylation to diverse transcript-selective regulatory networks.

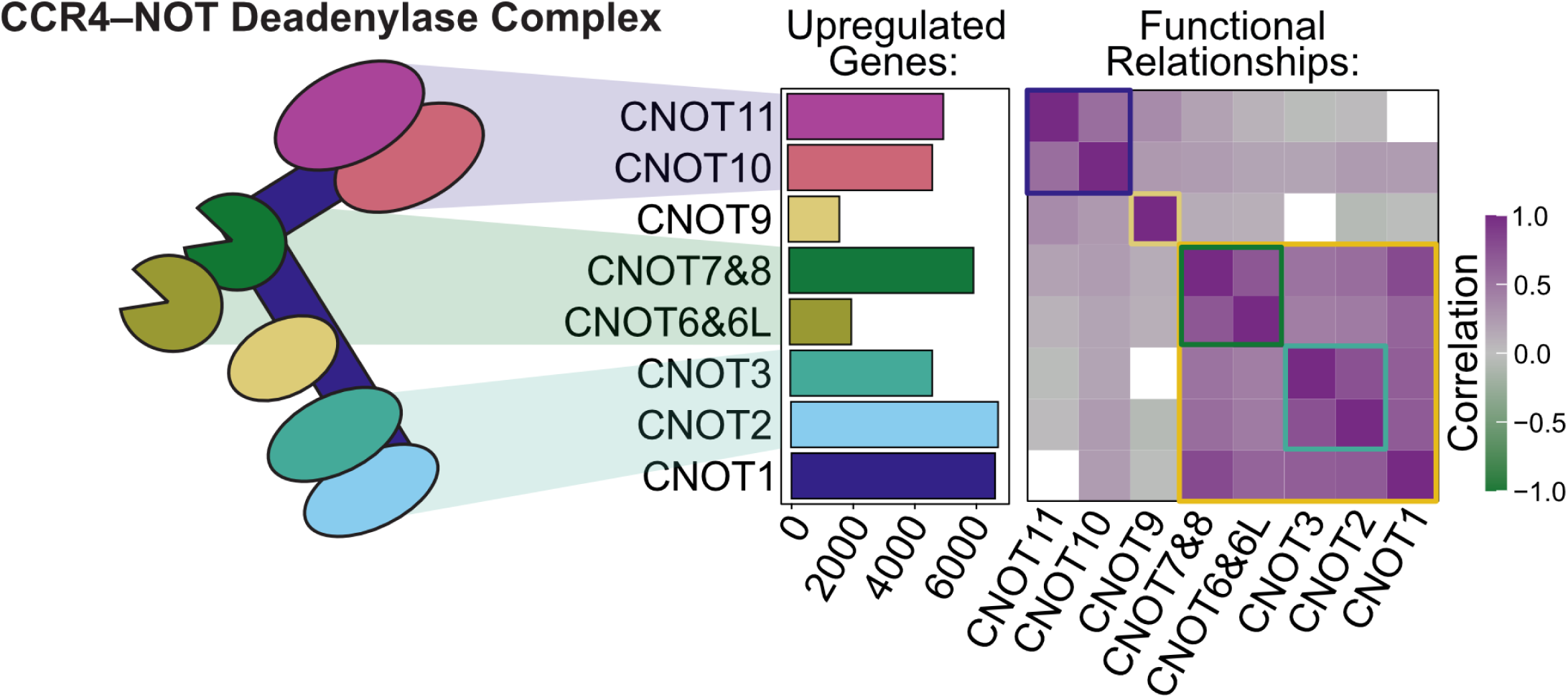

## Introduction

Messenger RNA decay plays a central role in the flow of genetic information by shaping the abundance, timing, and localization of gene expression. Decay rates differ widely among transcripts and can change in response to developmental cues, environmental changes, and cellular signaling, allowing rapid reprogramming of gene expression programs. Dynamic regulation of mRNA turnover is essential for processes such as cell cycle progression (1), cellular differentiation (2–4), immune responses (5), and embryonic development (3, 6–8). mRNA degradation also enforces quality control of gene expression by eliminating aberrant mRNAs (9, 10). In each of these contexts, mRNA degradation is highly regulated and selectively targeted.

mRNA degradation is catalyzed by specialized enzymes that act through coordinated pathways (9, 11–14). For most mRNAs, degradation begins with shortening of the 3′ poly(A) tail, a process known as deadenylation (9, 11–14). Once deadenylated, the mRNA is degraded by either of two pathways. In the 5′ to 3′ pathway, the 7-methyl-guanosine cap is removed by the decapping enzyme complex, allowing the XRN1 exoribonuclease to degrade the mRNA from the 5′ end (9, 11–14). Alternatively, the deadenylated mRNA can be degraded in a 3′ to 5′ direction by the exosome complex (9, 11–14). Although degradation of specific transcripts can be initiated by endonucleolytic cleavage, this mechanism appears to be less common (15, 16). Consequently, deadenylation-dependent decay is considered the main pathway of mRNA turnover and a nexus of post-transcriptional gene regulation.

Multiple deadenylase enzymes have been identified in eukaryotes (12). Among these, the evolutionarily conserved CCR4–NOT complex serves as the primary cytoplasmic deadenylase for mRNAs (17). Disruption of CCR4–NOT activity stabilizes mRNAs and increases poly(A) tail length (18–25). The human CCR4–NOT complex consists of eight core subunits organized into distinct biochemical modules **(Fig. 1A)** (18, 21). Although the structure of the intact complex has not yet been determined, high resolution structures are available for the four modules (25–34). The large CNOT1 protein forms the central scaffold upon which the complex is assembled. At its C-terminus, CNOT1 binds the CNOT2–CNOT3 heterodimer, forming the NOT module. The N-terminus of CNOT1 associates with the NOT10–11 module containing the CNOT10 and CNOT11 subunits, while the central region interacts with CNOT9 and the catalytic module.

**Figure 1.**
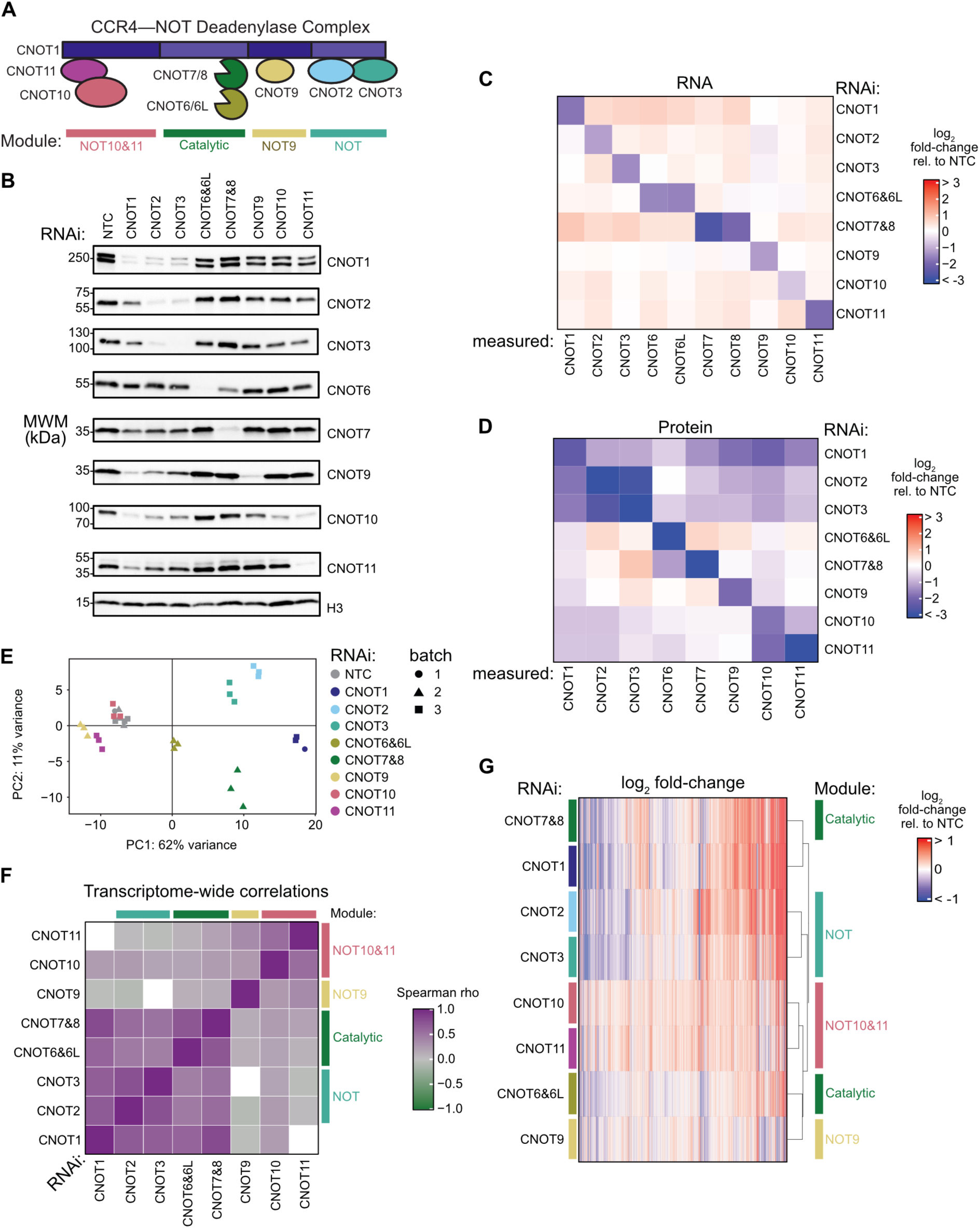
Comprehensive measurement of effects of CCR4–NOT on the transcriptome. (**A**) Diagram of the CCR4–NOT deadenylase complex subunits and biochemically defined structural modules. (**B**) Western blot validation of siRNA-mediated knockdown for each CCR4–NOT subunit in one of three replicate samples per knockdown condition. Western blots for the remaining two replicates are reported in **Figure S1**. Histone 3 (H3) served as a loading control. (**C**) Heat map of log_2_ fold-changes in the transcript abundance for each subunit in each knockdown condition, n=3. (**D**) Corresponding log_2_ fold-changes in protein abundance of each CCR4–NOT subunit in each knockdown condition measured by western blot, n=3. (**E**) Principal component analysis of RNA-seq samples for each subunit knockdown with three replicates per condition. Colors denote the targeted subunit or non-targeting control (NTC), and the data point shape indicates the batch in which the samples were collected (3 total batches, collected on 3 separate days). (**F**) Pairwise correlation matrix comparing transcriptome changes between knockdown conditions, showing Spearman rho correlation coefficients calculated from significance-weighted log_2_ fold-changes in transcript levels measured by RNA-seq. (**G**) Hierarchical clustering of the differential gene expression across all CCR4–NOT subunit knockdowns.

Despite detailed knowledge of its overall architecture, the molecular functions of several CCR4– NOT subunits and their specific contributions to gene regulation remain to be defined.

The catalytic module of CCR4–NOT contains a heterodimer of two classes of deadenylase: POP2–type and CCR4–type enzymes. In vertebrates, each class is represented by two paralogs with mutually exclusive incorporation in the complex: the POP2–type CNOT7 or CNOT8 (CNOT7/8) and the CCR4–type CNOT6 or CNOT6L (CNOT6/6L). The presence of two distinct deadenylase classes raises an important question: do these enzymes perform redundant or specialized functions? In cells, depletion-and-rescue experiments suggested that CNOT7/8 and CNOT6/6L have overlapping roles in bulk deadenylation (18). However, in vitro analysis with recombinant enzymes demonstrated that the two deadenylase classes generate distinct deadenylation products and differ in their ability to overcome poly(A)-binding protein mediated protection of the poly(A) tail (18, 35). Biochemical analysis of reconstituted CCR4– NOT complexes showed that assembly of the deadenylase heterodimer with the intact complex enhances catalytic activity (35), indicating that the non-catalytic subunits contribute to efficient deadenylation. Nevertheless, the specific roles of these non-catalytic subunits in mRNA deadenylation and decay remain incompletely understood in the cellular context.

In principle, CCR4–NOT can act on any poly-adenylated transcript; however, substantial evidence indicates that it preferentially engages specific RNAs through cis-acting determinants. These determinants can be recognized by sequence specific RNA-binding proteins and other recruitment factors that direct CCR4–NOT to target mRNAs, thereby accelerating deadenylation and decay (17, 21). Accumulating evidence further indicates that recruitment factors engage distinct subunits of the CCR4–NOT complex, which function as molecular adapters for regulatory inputs (17, 21). For example, the Pumilio family bind to specific sequences in target mRNAs and recruit CCR4–NOT through interactions with the NOT module, promoting deadenylation and decay of their targets (22, 36–40). Alternatively, CCR4–NOT can be recruited by the microRNA induced silencing complex (miRISC), in which the TNRC6A/B/C proteins interact with the CNOT1 and CNOT9 subunits to stimulate deadenylation and repression of microRNA targets (21, 41–45).

The repertoire of CCR4–NOT recruitment factors continues to expand (17, 21). Given that the human genome encodes more than 1, 500 RNA-binding proteins and approximately 2000 microRNAs, the potential diversity of mechanisms that regulate CCR4–NOT recruitment is likely large (46, 47). A major challenge for the field is to comprehensively define how these regulatory inputs control CCR4–NOT mediated deadenylation and decay across the transcriptome.

Selective deadenylation by CCR4–NOT can also be directed through translation-dependent mechanisms. In the quality control pathway Nonsense-Mediated Decay, NMD factors can recruit CCR4–NOT to accelerate degradation of transcripts containing premature termination codons (48, 49). CCR4–NOT is also involved in codon optimality mediated mRNA decay, in which it is recruited to ribosomes stalled at non-optimal synonymous codons, leading to selective degradation of the mRNA (50–53). More recently, CCR4–NOT was shown to be selectively recruited during decoding of specific arginine codons through a mechanism termed P-site tRNA-mediated decay (54). In another specialized pathway, the nascent peptide encoded by tubulin mRNAs recruits CCR4–NOT to maintain tubulin homeostasis. This mechanism is mediated by TTC5 and SCAPER, which bind the NOT10–11 module (55). These examples demonstrate that CCR4–NOT can integrate diverse signals arising during translation, including premature termination, codon usage, tRNA decoding, and nascent peptide recognition, to selectively regulate mRNA stability. However, the role of CCR4–NOT in these translation-dependent processes has not been comprehensively analyzed.

In this study, we investigated the global impact of each CCR4–NOT subunit in mRNA deadenylation and decay in human cells. High throughput RNA sequencing (RNA-seq) and poly(A) length analysis revealed that CCR4–NOT regulates transcripts from thousands of genes, many of which were shown to be physically associated targets using a crosslinking-based capture approach. Individual subunits contributed to the selective decay of shared and unique pools of mRNAs, and their effects largely reflected the biochemically defined modular architecture of the complex. The CNOT7 and CNOT8 deadenylases were primary drivers of deadenylation and decay, whereas the CNOT6 and CNOT6L regulated a more limited subset of targets. Bioinformatic analysis identified processes and pathways that are regulated by CCR4– NOT and uncovered new candidate RNA determinants and recruitment factors that contribute to target specificity. Collectively, these findings expand our understanding of CCR4–NOT mediated gene regulation and provide a resource for dissecting its mechanisms and biological functions.

## Materials and Methods

### Cell Culture

Human HCT116 cells (ATCC, CCL-247) were cultured in McCoy’s 5A modified medium (Thermo Fisher Scientific, 16600082) supplemented with 10% (v/v) fetal bovine serum (Genesee Scientific, 25-514) and antibiotics (100 U/mL penicillin and 100 µg/mL streptomycin, Thermo Fisher Scientific), at 37 °C with 5% CO_2_.

### siRNA transfection

Catalog numbers and sequences for siRNAs are listed in **Supplementary File S1**. HCT116 cells were seeded at 200, 000 cells in 2 mL medium per well in a 6-well plate (USA Scientific, CC7682-7506). After 24 hours, the cells were transfected using Dharmafect 4 (Horizon Discovery) with 25 nM final concentration of On-Target Plus Smartpool siRNAs (Horizon Discovery) for the indicated protein of interest. A second transfection of siRNA was performed after 24 hours for the CNOT7 and CNOT8 (combined knockdown), CNOT6 and 6L (combined knockdown), CNOT10, and CNOT11 knockdown conditions. 72 hours after the initial siRNA treatment, cells were washed with phosphate-buffered saline (PBS), pH 7.4 (Thermo Fisher Scientific) and collected by scraping followed by centrifugation at 200 x g for 5 mins. Three biological replicates were collected for each knockdown condition with a matched non-targeting control (NTC) siRNA sample. RNAi depletion was optimized to maximize depletion of each subunit (**Supplementary File S2**).

For cell viability measurements under RNAi knockdown conditions, HCT116 cells were counted and assessed for cell viability by trypan blue staining (Thermo Fisher, T10282) using the Countess Automated Cell Counter (Thermo Fisher). These measurements were taken from three biological replicates and two technical replicates with a total of six replicates per condition, with the exception of CNOT1, which had four total replicates.

### RNA purification

Total RNA was extracted from the collected cell pellets using the SimplyRNA cells kit and Maxwell RSC instrument (Promega) following the manufacturer’s protocol. Twice the amount of DNase was added for the on-bead DNase treatment. Purified RNA was eluted in 40 μL of nuclease-free water and quantified using a Nanodrop spectrophotometer (Thermo Fisher Scientific). The quality of the RNA was determined by Agilent Tapestation analysis with RNA integrity number (RIN) values ≥9.9 for all samples.

### RNA-seq library preparation and sequencing

Total RNA was purified as described above from three biological replicate samples per knockdown condition. External RNA Controls Consortium (ERCC) Spike-In Control Mixes were included for normalization in all samples at 2 µL of a 1:1000 dilution for 100 ng total RNA input according to manufacturer’s protocol (Thermo Fisher Scientific, 4456740). RNA-seq libraries were prepared using the Stranded Total RNA Library Prep with RiboZero Plus (Illumina) and sequenced with paired-end, 150 bp reads on a NovaSeq S4 (Illumina) with approximately 81 million mean read-pairs per sample. The library preparation and RNA sequencing was performed by the University of Minnesota Genomics Center. RNA-seq was performed according to the standard ENCODE guidelines reported in **Supplementary File S3**. Reads were uploaded to GEO under accession number GSE320579.

### RNA-Seq Processing and Differential Expression Analysis

RNA-seq data were processed using the Nextflow RNA-seq pipeline (nf-core/rnaseq v3.9; NXF_VER 21.10.3) with Singularity (v3.1.1) (56, 57). The UCSC hg38 genome FASTA was combined with ERCC92 spike-in sequences for the -fasta input, and a GTF file combining hg38 UCSC refGene annotations with ERCC92 annotations was used as the -gtf input. Normalized counts were generated with DESeq2 (v1.44.0; estimateSizeFactors) using ERCC92 spike-in genes as the control set, and batch effects were removed with limma (v3.58.1; removeBatchEffect). Principal component analysis (PCA) was performed on variance-stabilized data (vsd) using the top 500 most variable genes (DESeq2, v1.44.0; plotPCA).

The 16, 547 genes with at least 10 counts in at least three samples were retained for differential expression analysis by DESeq2 (v1.44.0). Significantly up or downregulated genes were determined using Benjamini–Hochberg adjusted p-values (p^adj^ < 0.05) and no log_2_ fold-change cutoff. The resulting DESeq2 outputs are presented in **Supplementary File S4**. For analysis of PUM1&2 RNAi, DESeq2 was repeated using PUM1&2 RNAi samples and all CCR4–NOT conditions.

To visualize differential expression results, MA plots were generated for each knockdown versus NTC control, with log_2_ fold-changes shrunken using the apeglm method (DESeq2, v1.44.0) to reduce noise from low-count genes. Base mean expression values were plotted against shrunken fold-changes, with significant genes (p^adj^ < 0.05) highlighted. Heat maps of log_2_ fold-changes across conditions were generated with ComplexHeatmap (v2.20.0), and organized by hierarchical clustering using Euclidean distance and complete linkage based on log_2_ fold-change values. Overlaps of significantly upregulated genes were visualized using Venn diagrams (euler, v7.0.2).

### Functional Enrichment Analyses

Gene ontology (GO) enrichment analysis was performed with clusterProfiler (v4.12.0; enrichGO), restricted to biological process (BP) terms, with top terms selected by adjusted p-value. Gene set enrichment analysis (GSEA) was carried out using clusterProfiler (v4.18.4; gseGO). Genes were ranked in descending order by the DESeq2 Wald test statistic. To identify enriched GO terms, GSEA was conducted for BP terms from org.Hs.eg.db (v3.22.0), with enrichment tested via built-in permutation tests using default parameters. The full results of both analyses are presented in **Supplementary File S5**. Curated gene sets were retrieved from MSigDB via msigdbr (v7.5.1) (58) for hallmarks of apoptosis, DNA repair pathways, and tubulin genes, or from selected publications for: cell cycle phases (59), hallmarks of cancer (60), and nonsense-mediated mRNA decay (NMD) targeted gene sets (61). For curated datasets, GSEA was conducted using clusterProfiler (v4.12.0) and fgsea (v1.26.0). Enrichment was tested with fgsea (nperm = 10, 000). Normalized enrichment scores (NES) and adjusted p-values are reported in **Supplementary File S5**.

### RNA Binding Protein Analysis

Previously compiled CLIP-seq datasets from CLIPdb 1.0 were used to infer RBP–target interactions, focusing on binding events located within 3′ UTRs (62, 63). The mean log_2_ fold-change of each RBP target in each CCR4–NOT condition was calculated and significance was determined relative to genes that are not targets of that RBP. Comparisons with known protein-protein interactions were compiled from: (24, 64). Subcellular localization (cytoplasmic or nuclear) was curated from UniProt (65).

### Motif analysis via FIRE

Enriched motifs were identified using the FIRE algorithm (66). Genes expressed in all datasets were associated with representative transcripts and binned by the DESeq2 Wald statistic. FIRE identifies motifs relatively enriched or depleted in the 3′UTRs of these representative sequences. The outputs of FIRE are presented in **Supplementary File S6** and were converted to genomic coordinates for downstream analysis. Overlap between identified motifs was quantified via the Jaccard statistic.

### Nanopore library preparation and sequencing

Poly(A) RNA was enriched using the Poly(A) RNA Selection Kit (Lexogen) according to the manufacturer’s instructions. Direct RNA sequencing libraries were prepared from poly(A) RNA using the SQK-RNA002 kit (Oxford Nanopore Technologies, Oxford, UK) and sequenced on a GridION using FLO-MIN106D flow cells, one flow cell per library. Two biological replicates were analyzed for each knockdown condition using independent flow cells and four replicates were performed for the non-targeting control.

For each flow cell, reads passing the real-time (MinKNOW) quality filter applied during sequencing (fast5_pass) were converted and consolidated into a single pod5 file using the pod5 package (v0.3.10). Reads were basecalled on GPU compute nodes with Dorado (v0.7.0) using the rna002_70bps_hac@v3 model with the --estimate-poly-a flag, which records a per-read poly(A) tail length estimate in the pt:i tag. In the same step, reads were aligned to the GENCODE v38 human transcriptome (67) using Dorado’s integrated minimap2 aligner (68), and the resulting BAM file was coordinate-sorted and indexed with samtools (69). Poly(A) tail lengths were extracted from the pt:i tags, and reads for which a tail length could not be estimated (pt values of -1 or 0) were discarded. Transcript-level abundances were estimated with NanoCount (70), which uses an expectation-maximization algorithm to probabilistically assign multi-mapping long reads. Raw direct RNA sequencing data was uploaded to GEO under accession number GSE320579.

Bulk tail length distributions were summarized independently for each flowcell. All poly(A) measurements for each sample are presented in **Supplementary File S7**, with summary statistics (mode, median, mean, standard deviation, and interquartile range) presented in **Supplementary File S8**. For gene-level poly(A) analysis, all poly(A) measurements and associated genes, for all flowcells, were combined and limited to the 6, 951 genes represented by at least 10 poly(A) measurements in all flowcells. Mean, steady-state poly(A) measurements in control and CNOT7&8 RNAi samples were calculated using the R package lme4 (v2.0.1), which considered the in-sample variation and variation between replicates. Comparisons between steady-state poly(A) measurements and the effect of CNOT7&8 RNAi were conducted on the 6, 546 genes that could be unambiguously associated with differential gene expression analysis.

### CNOT7 crosslinking and RNA immunoprecipitation

Formaldehyde-crosslinked RNA immunoprecipitation was performed as previously described (71), using a dox-inducible V5-tagged CNOT7 catalytic mutant (D40A; V5-CNOT7^MUT^) system engineered in HCT116 cells. The cells were crosslinked with 0.1% formaldehyde for 10 mins, lysed, and subjected to V5 immunoprecipitation. Following crosslink reversal, co-purifying RNAs were isolated, and RNA-seq libraries were generated from three biological replicates of HCT116 cells, using V5-CNOT7^MUT^-immunoprecipitated RNA from dox-induced cells with matched input controls. Ribosomal RNA depletion was performed prior to library preparation. Sequencing was performed using single-end 75 bp reads on a NextSeq500 platform (Genomics Platform at the Institute for Research and Immunology and Cancer, Montreal). Raw read quality was assessed using FASTQC v0.12.1. Reads were aligned to the GRCh38 reference genome using STAR and raw counts were calculated with FeatureCounts (release 110). RNAs were considered significantly enriched if they exhibited >1.5-fold enrichment relative to input with an Benjamini– Hochberg adjusted p-value (p^adj^) of <0.01 relative to input. Raw RIP-seq data was uploaded to GEO under accession number GSE348528. The processed results are reported in **Supplementary File S9**.

### Western blotting

Antibodies used in this study are listed in **Supplementary File S1**. Cell pellets were resuspended in 100 µL of RIPA buffer (25 mM Tris–HCl pH 7.6, 150 mM NaCl, 1% NP-40, 1% sodium deoxycholate, 0.1% SDS) supplemented with 2x cOmplete^TM^ Protease Inhibitor Cocktail (Roche) and lysed by cell disruption. Total protein concentration was quantified using DC-Lowry assay (BioRad). 10 µg of each protein sample was added to SDS-PAGE loading dye (80 mM Tris-HCl pH 6.8, 10% glycerol, 2% SDS, 100 mM DTT, 0.2% bromophenol blue, 100 mM 2-Mercaptoethanol), heated at 85 °C for 10 mins, and loaded onto a 4-20% gradient SDS polyacrylamide gel (Bio-Rad) and run at 195V for 45 mins. The proteins were transferred onto a PVDF membrane (Immobilon P, Millipore) at 60V constant for 1.5 hours at 4 °C. The membrane was blocked for 1 hour with 5% w/v nonfat powdered dry milk, 1X PBS, and 0.1% Tween-20 (PBST). The indicated primary antibody was diluted 1:1, 000 in PBST and rocked overnight at 4 °C. The next day, the membrane was washed 3 times 10 mins in PBST. The appropriate secondary antibody HRP was diluted 1:10, 000 in PBST and added to the membrane rocking at RT for 1 hour. The membrane was washed 3 times 10 mins in PBST. The washed membrane was incubated with Immobilon Western Chemiluminescent HRP Substrate (Millipore) for 1 min and visualized on an Azure 300 (Azure Biosystems). Band intensities were quantified using Image Lab software (Bio-Rad).

### Plasmids and Cloning

The oligonucleotides and plasmids used in this study are listed in **Supplementary File S1**. The Nano-luciferase (Nluc) reporter empty vector (pNLP MCS) was generated as previously described (22). The firefly luciferase (Fluc) plasmid pGL4.13 (Promega) was used as a transfection efficiency control. The Nluc 3xCNOT ETS1 (3xCM1) and 6xCNOT ETS1 (6xCM1) reporter plasmids were created by inserting three or six copies, respectively, of a sequence containing a composite enriched motif (5′-tgtcgatgTTTTTTTAAAA-3′) for CNOT1, CNOT10, and CNOT11, which was identified using FIRE within the ETS1 3′UTR (Gene ID: 2113), into the Nluc 3′UTR by oligo cloning.

### Luciferase Reporter Assays

HCT116 cells were seeded with 5, 000 cells in 100 μL of growth medium per well in a 96-well white-walled plate (Fisher, 12-566-00). After 24 hours, the cells were transfected with CNOT1, CPEB4, or non-targeting control siRNAs. 24 hours after siRNA treatment, the cells were transfected with 5 ng Fluc reporter plasmid, 10 ng of Nluc reporter plasmid, and 85 ng pF5A empty vector control (100 ng total transfected DNA per well). The transfected cells were grown for another 48 hours and luciferase values were measured using the Dual-Glo Assay system (Promega) and a GloMax Discover luminometer. The RNAi knockdown efficiency was verified by western blot analysis for CNOT1 depletion and RT-qPCR for CPEB4 depletion.

To analyze reporter data, a relative response ratio (RRR) was calculated by normalizing the Nluc luciferase activity (measured in relative light units, RLU) against the corresponding Fluc RLU for each sample. These RRR values were then used to calculate the fold-change between the Nluc the 3x CNOT motif (3xCM1) and 6x CNOT motif (6xCM1) reporters and the Nluc empty vector (multiple cloning site, MCS) to measure repression of expression. All reporter assays were performed in three independent experiments, each of which had 3 biological replicates, for a total of nine replicates. The data is reported as mean log_2_ fold-change with the standard deviation. The statistical significance of comparisons was calculated using an ordinary one-way ANOVA and Tukey test for multiple comparisons.

### Flow Cytometry Cell Cycle Analysis

HCT116 were seeded and transfected with siRNAs as described above. 72 hours after the initial siRNA treatment, cells were washed with phosphate buffered saline (PBS) pH 7.4 (Thermo Fisher Scientific) and trypsinized using TrypLE Express (Thermo Fisher Scientific, 12605028).

The cells were counted using trypan blue staining (Thermo Fisher) and the Countess Automated Cell Counter (Thermo Fisher). 4 million cells per condition were collected by centrifugation at 200 x g for 5 mins. The cell pellets were suspended in 200 µL cold PBS and vortexed to mix. Cells were added dropwise to 4 mL ice cold 70% ethanol (Fisher Scientific) in a 15 mL conical centrifuge and vortexed to mix. The mixture was incubated for overnight fixation at -20 °C. The next day, cells were collected by centrifugation at 1, 200 rpm for 10 min at 4 °C. The supernatant was removed and cells suspended in 1 mL of a final concentration of 40 μg/mL propidium iodide (Invitrogen), 100 μg/mL RNase A (Promega), and PBS. The mixture was incubated at 37 °C for 30 mins prior to analysis by flow cytometry using a BD LSRFortessa (BD Biosciences).

### Reverse transcription and quantitative polymerase chain reaction

The RTqPCR parameters, including the primer sequences, amplification efficiencies, and amplicon sizes, following the standard MIQE guidelines are reported in **Supplementary File S1**. RNAi-mediated knockdown of CPEB4 versus the non-targeting control (NTC) was measured by RTqPCR. RNA was isolated using the Maxwell system as described above. 1 µg of RNA was reverse transcribed with the LunaScript RT kit (New England Biolabs) and 1 µL of random hexamer oligonucleotides (Promega) per reaction. Reverse transcription was performed at 25 °C for 2 min, 55 °C for 10 min, and 95 °C for 1 min. Quantitative PCR was performed using 50 ng of cDNA for CPEB4 and 0.5 ng of cDNA for 18S rRNA control. Each reaction contained 0.25 µM of each primer and Luna Universal qPCR Master Mix (New England Biolabs), following the manufacturer’s protocol. As negative control, a no-RT reaction was analyzed for each primer set. The cycling conditions were: (i) 95 °C for 2 min, (ii) 95 °C for 15 sec, and (iii) 60 °C for 1 min. Steps ii–iii were repeated for a total of 40 cycles. A melt curve was generated over a range of 65 °C to 95 °C. Products were analyzed by gel electrophoresis to verify the correct amplicon size. Quantification cycle (Cq) values were obtained using CFX Manager software (Bio-Rad).

Relative expression was calculated using the Livak method, with normalization to 18S rRNA.

### Viability and Apoptosis Assay

The ApoLive-Glo Multiplex Assay (Promega, G6410) was performed according to the manufacturer’s instructions. HCT116 cells were seeded at 200, 000 cells in 2 mL complete growth medium per well in a 6-well plate. For RNAi-mediated depletion, the cells were treated with CNOT1, CNOT6&6L, CNOT7&8, and NTC siRNAs, as described above. 72 hours after the initial siRNA treatment, cells were washed with PBS pH 7.4 and trypsinized using TrypLE. The cells were counted using trypan blue staining and the Countess Automated Cell Counter and reseeded at 30, 000 cells per well in 100 μL of prewarmed complete growth media into a white-walled 96-well plate (Fisher, 12-566-00). 20 µl of viability reagent was added to each well and the cells were incubated at 37 °C for 30 mins. Fluorescence was then measured at 400 nm excitation and 505 nm emission on a GloMax Discover luminometer (Promega) to assess cell viability. Viability is determined by a live-cell protease activity within intact cells which produces fluorescent signals proportional to the number of living cells. After the viability assay, 100 µl of Caspase-Glo 3/7 reagent was added to all wells and incubated at room temperature for 30 mins. Luminescence was then measured (GloMax Discover luminometer, Promega) to detect caspase-3/7 activity, a key biomarker of apoptosis.

### Mitochondrial Toxicity Assay

The mitochondrial ToxGlo assay (Promega, G8000) was performed according to the manufacturer’s instructions. HCT116 cells were seeded at 200, 000 cells in 2 mL complete growth medium per well in a 6-well plate. For RNAi-mediated depletion of each CCR4–NOT subunit, the cells were treated with siRNAs as described above. 72 hours after the initial siRNA treatment, cells were washed with PBS pH 7.4 and trypsinized using TrypLE. The cells were counted using trypan blue staining and the Countess Automated Cell Counter and 200, 000 cells per condition were collected. Cells were washed twice with PBS pH 7.4, suspended in DMEM growth medium without glucose and without FBS (Thermo, 11966025) supplemented with 10 mM final concentration of galactose (Sigma, G5388). Cells were cultured in medium with galactose as a sole sugar source to increase the assay sensitivity according to the manufacturer’s protocol. Cells in DMEM + galactose were re-seeded at 10, 000 cells per well in triplicate. As a positive control for membrane integrity, cells were treated with 10 μL of a 0.8 mg/ml working stock of digitonin (Promega, G9441). As a positive control for mitochondrial toxicity, cells were treated with a final concentration of 3 μM of the mitochondrial toxin antimycin A (Sigma, A8674). DMSO at equivalent volume was used as a vehicle-only negative control.

Cells were incubated at 37 °C for 90 mins and then 20 μL of 5x cytotoxicity reagent was added to each well and incubated for an additional 30 mins at 37 °C. Fluorescence was measured at 485 nm excitation and 500–550 nm emission on a GloMax Discover luminometer (Promega), to first assess cell membrane integrity. The plate was then equilibrated to room temperature for 10 mins. 100 μL of the ATP detection reagent was added and incubated at room temperature for 5 mins before measuring luminescence to measure the amount of ATP produced by oxidative phosphorylation. The percent response was calculated by dividing measured value for ATP (luminescence) by the untreated control for each condition using GraphPad Prism.

### Statistical Analysis

All analyses were performed in R. Unless otherwise specified, plots were generated using the R package *ggplot2* and related packages *ggprism*, *ggpointdensity*, *ggrepel*, *ggridges*, *ggseqlogo*, *ggbreak* and *viridis* as needed. We compared the distributions of the following metrics between defined gene sets: translation efficiency (72), optimal codons based on codon stability coefficient (CSC) values (73), fraction of non-optimal arginine codons (54), and mRNA half-lives (74). All data used for these comparisons is presented in **Supplementary File S5**. For comparing the distributions of these metrics, only the genes with measurements in all RNAi conditions were considered. These distributions were displayed via boxplots, showing medians with quartiles, with whiskers extending to 1.5X interquartile range unless otherwise noted. The significance of the differences between these distributions was determined via a two-sided Wilcoxon rank-sum test performed using the package *ggpubr*, which interfaces with package *rstatix*, the package *stats*, and base R functions. Reported p-values are specified in the figure legends. The magnitude of differences between distributions was assessed via Cliff’s delta (δ) from the package *effsize,* and is noted within the figures. For poly(A) analysis, δ was calculated based on the frequency of the poly(A) measurements using base R functions. Following standard convention, we described differences in distributions as negligible (|δ| < 0.147), small (0.147 ≤ |δ| < 0.33), and large (|δ| ≥ 0.474). Summary statistics for the distributions are presented in **Supplementary File S5**.

When comparing overlaps in Venn diagrams, the significance of overlap was determined via a Fisher’s exact test with adjusted p-value. The number of genes in each class, odds ratio, and adjusted p-values are presented **Supplementary File S5**. Fisher’s Exact Tests were used to determine whether motifs were likely to occur within the 3′UTRs of genes regulated by other CCR4–NOT subunits.

## Results

### Comprehensive measurement of effects of CCR4–NOT on the transcriptome

To define the contribution of individual CCR4–NOT subunits to transcriptome regulation, we depleted each of the ten subunit proteins across eight RNAi knockdown conditions (**Supplementary File S2**) and measured changes in RNA abundance by RNA-seq relative to cells transfected with non-targeting control siRNAs (NTC). Previous studies demonstrated functional overlap between paralogs, so CNOT6 and CNOT6L were depleted together (CNOT6&6L), as were CNOT7 and CNOT8 (CNOT7&8) (18, 64). HCT116 cells were used because they possess a near-diploid karyotype, retain an intact p53 pathway, and support efficient RNAi. Knockdown conditions were optimized for depletion of each CCR4–NOT protein (**Fig. 1B**) without loss of cell viability (**Fig. S1A**) (22). To assess reproducibility, three biological replicates were measured for each knockdown. The RNA samples were collected at 72 hours post-siRNA transfection in three experimental batches and spike-in controls were included for normalization. Ribosomal RNA-depleted libraries were prepared from purified RNA from each sample and then were sequenced by Illumina RNA-seq, yielding approximately 81 million read-pairs each.

From the RNA-seq data, depletion of mRNA encoding each CCR4–NOT subunit was observed (**Fig. 1C**). Depletion of each CCR4–NOT subunit was confirmed by western blots from the three biological replicates (**Fig. 1B and S1B**) and quantitated (**Fig. 1D, Supplementary File S2**).

Consistent with previously reported stabilizing interactions within CCR4–NOT, depletion of specific subunits altered the abundance of their binding partners (**Fig. 1D**). Depletion of CNOT1, upon which the complex is scaffolded, reduced protein abundance of all subunits.

Interdependence of protein levels was also observed for the components of the NOT and NOT10–11 modules, allowing us to assess the functional role of each module.

Principal component analysis separated samples primarily according to knockdown condition rather than experimental batch, with biological replicates clustering tightly (**Fig. 1E**). CNOT1 depletion accounted for the greatest variance, consistent with its role as the scaffold of the complex, whereas the remaining subunits distributed according to their established modular organization. Samples in which the NOT module components CNOT2 and CNOT3 were depleted clustered closely. Likewise, samples in which the catalytic subunits CNOT6&6L and CNOT7&8 were depleted occupied neighboring but distinct positions, consistent with their related enzymatic functions. CNOT9, CNOT10, and CNOT11 samples were clearly separated from the NOT and catalytic modules, and clustered near the NTC condition. These results suggest that CCR4–NOT modules regulate distinct networks of transcripts.

### Transcriptome responses reveal that function follows the modular form of CCR4–NOT

We measured changes in gene expression upon depletion of the CCR4–NOT subunits, calculated relative to NTC condition, with an adjusted p-value of <0.05 (**Fig S2A-B**). Knockdown of each subunit caused significant changes in expression levels of many genes. Depletion of CNOT1, the NOT module, and CNOT7&8 deadenylases had the largest effects in terms of number of genes (**Fig. S2A-B**) and magnitude of changes (**Fig. S3A**).

To assess functional relationships among CCR4–NOT subunits, we examined the correlations between the transcriptome-wide changes in RNA abundance in response to each knockdown using Spearman’s rank correlation coefficient (**Fig. 1F**). Knockdowns of subunits within the same module produced the most similar gene expression changes. Strong correlations were observed between members of the NOT and catalytic modules (Spearman rho 0.44 to 0.80), as well as the NOT10–11 module (0.55). In contrast, depletion of CNOT9 showed relatively weak correlations with the other knockdowns (-0.06 to 0.12), suggesting that it regulates a largely distinct set of RNA targets. These relationships are further supported by hierarchical clustering of the differentially expressed genes across all knockdown conditions (**Fig. 1G**), which largely grouped subunits according to their biochemically-defined modular organization. One notable exception involved the catalytic module: CNOT1 clustered most closely to CNOT7&8 rather than the other structural subunits. This likely reflects their dominant roles as the principal scaffold and deadenylase activities of the complex, respectively, and their broad effects on gene expression. In contrast, CNOT6&6L clustered with CNOT9, consistent with their comparatively modest impact on the transcriptome.

Because CCR4–NOT promotes mRNA decay, we next focused on transcripts that increased in abundance following depletion of individual subunits, reasoning that these RNAs are most likely to represent direct decay targets. Depletion of each subunit increased the abundance of thousands of mRNAs, with knockdown of CNOT1, the NOT module (through CNOT2), and CNOT7&8 affecting the largest number of genes (**Fig. 2A**). These results indicate a broad role of CCR4–NOT in mRNA decay.

**Figure 2.**
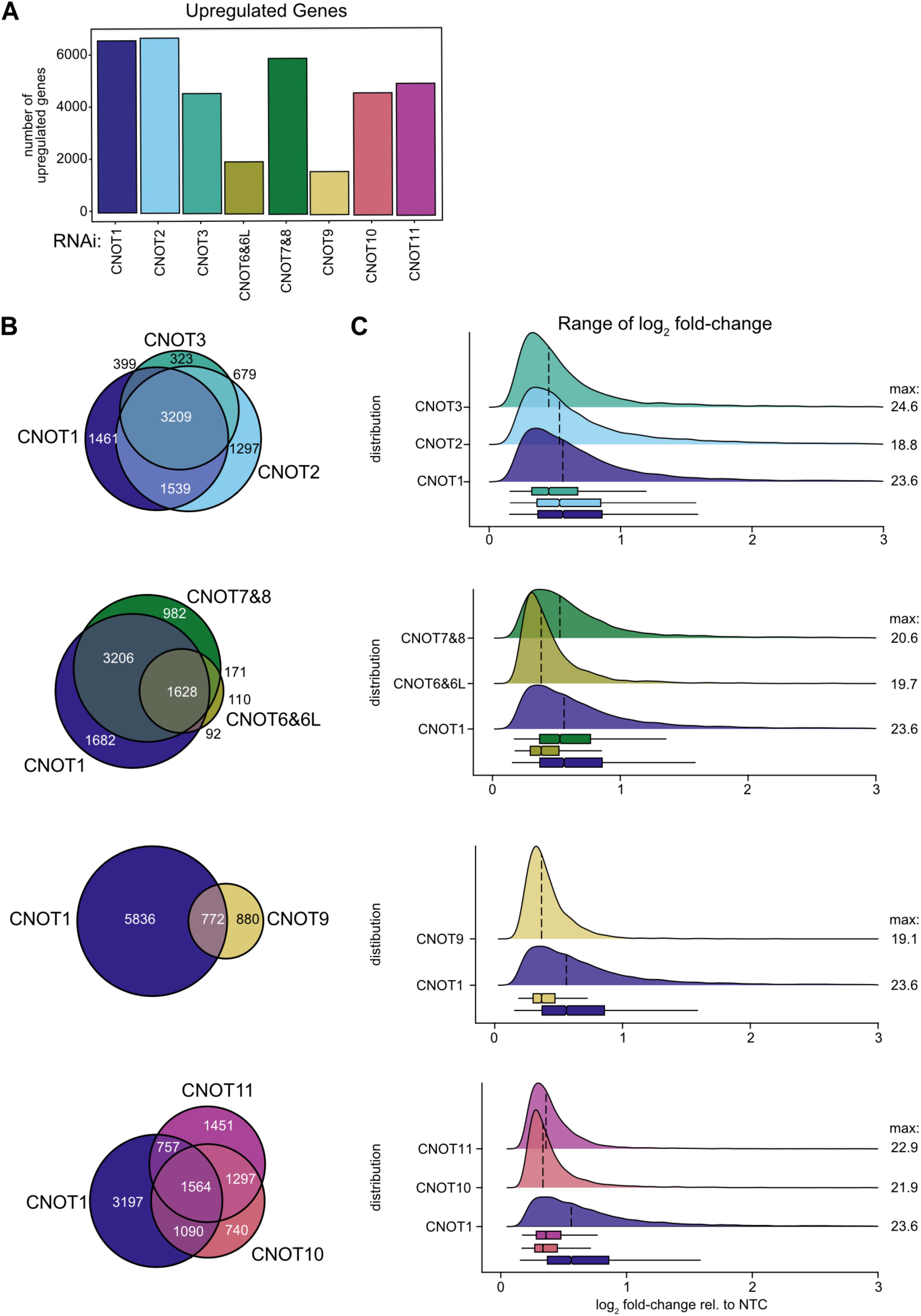
Relationship of transcriptome responses to modular architecture of CCR4–NOT subunits. (**A**) Bar graph comparing the number of genes upregulated in response to RNAi-mediated depletion of each CCR4–NOT subunit. (**B**) Venn diagrams illustrating the number of overlapping genes upregulated by each knockdown, grouped by CCR4–NOT module. All pairwise comparisons shown were significant based on Fisher’s exact test with p-values adjusted via the Benjamini-Hochberg method (see **Supplementary File S5**). (**C**) Ridgeline plots displaying distributions of log_2_ fold-changes of transcript abundance, as kernel density estimates, for genes upregulated by knockdown of each CCR4–NOT subunit. Plots are organized by module. The black bar marks the median of each distribution, and corresponding boxplots are below. For all plots, the window was limited to 0 ≥ log_2_ fold-change ≥ 3 for clarity, and the maximum log_2_ fold-change observed for each dataset is listed on the right. Full distributions presented in **Fig. S3A**.

To assess functional relationships among CCR4–NOT modules, we compared both the overlap of upregulated genes (**Fig. 2B, Fig. S3B, Supplementary File S5**) and the magnitude of their changes (**Fig. 2C**) compared to CNOT1 knockdown. The transcriptome-wide similarities observed between the NOT and catalytic modules (**Fig. 1F**) were preserved within the population of upregulated transcripts (**Fig. 2B**). Depletion of CNOT1 and the NOT module components CNOT2 and CNOT3 resulted in highly similar sets of upregulated transcripts and comparable increases in mRNA abundance (**Fig. 2B, 2C**). Among genes uniquely upregulated by CNOT2 and CNOT3, similar responses were observed with depletion of other CCR4–NOT subunits (**Fig. S3C-E**), albeit changes were not to the level of significance. Given the interrelated stability of the NOT module (**Fig. 1D**), the relatively high number of genes uniquely regulated by CNOT2 was surprising. However, the distributions of log_2_ fold-changes for CNOT2 unique genes (**Fig. S3D**) and CNOT3 unique genes (**Fig. S3E**) show similar responses with CNOT3 and CNOT2 depletion, respectively, as well as with depletion of NOT10–11 module.

Furthermore, of the 623 genes uniquely regulated by CNOT2, 348 (56%) were increased by CNOT1 depletion (**Fig. S3C**). Similarly, of the 159 genes uniquely regulated by CNOT3, 90 (57%) were increased by CNOT1 depletion (**Fig. S3C**). Adding to the transcriptome-wide correlations and interdependent subunit stabilities, the overlap between upregulated genes suggests an interconnection of function between the NOT module, CNOT1 scaffold, and catalytic module.

Depletion of CNOT7&8 deadenylases increased many of the same transcripts affected by CNOT1 and to a similar extent (**Fig. 2B, 2C**). In contrast, depletion of CNOT6&6L affected fewer transcripts overall (**Fig. 2A**). These transcripts largely overlapped with those regulated by CNOT1 and CNOT7&8 (**Fig. 2B**) but exhibited more modest increases in abundance (**Fig. 2C**). Together, these findings indicate that CNOT1, the NOT module, and CNOT7&8 are the major effectors of CCR4–NOT mediated mRNA decay and that CNOT7&8 are the predominant deadenylases.

Among all subunits, depletion of CNOT9 resulted in the fewest upregulated genes (**Fig. 2A**) and the smallest changes in mRNA abundance (**Fig. 2C**). Consistent with its distinct behavior within the complex, including its independence in protein stability (**Fig. 1D**) and weak transcriptome-wide correlation with the other CCR4–NOT subunits (**Fig. 1F**), the overlap between transcripts regulated by CNOT9 and CNOT1 was lower than observed for other modules (**Fig. 2B**).

Together, these findings suggest that CNOT9 has a largely non-redundant role in CCR4–NOT mediated mRNA decay.

Depletion of CNOT10 or CNOT11 increased the abundance of thousands of transcripts (**Fig. 2A**), although the magnitudes of these changes were lower than CNOT1 depletion (**Fig. 2C**). Consistent with their mutual dependence for protein stability (**Fig. 1D**) and clustering in transcriptome-wide expression correlations (**Fig. 1F**), CNOT10 and CNOT11 affected overlapping sets of transcripts (**Fig. 2B**). However, their overlap with CNOT1-regulated transcripts was more limited than that observed for the NOT and catalytic modules (**Fig. 2B**). Notably, nearly half of the genes upregulated by CNOT10 or CNOT11 knockdown were not upregulated by CNOT1 depletion. Among all knockdown conditions, transcripts uniquely upregulated by depletion of CNOT11 or CNOT10 were largely unaffected or downregulated following depletion of other subunits (**Fig. S3C**). Of the 380 genes uniquely upregulated by CNOT10 knockdown, only 119 (31%) were increased by CNOT1 depletion. Of the 726 genes uniquely upregulated by CNOT11 knockdown, only 126 (17%) were increased by CNOT1 depletion. Similarly, the 1, 564 CNOT10 and CNOT11 co-targets that overlap with CNOT1 were affected by depletion of most CCR4–NOT subunits, while the 1, 297 genes that did not overlap with CNOT1 were largely unaffected or downregulated (**Fig. S3F**). The distributions of log_2_ fold-changes for CNOT10 unique genes (**Fig. S3G**) and CNOT11 unique genes (**Fig. S3H**) show similar responses with CNOT11 and CNOT10, respectively. These findings suggest that the NOT10–11 module regulates a distinct subset of transcripts and may carry out functions that are at least partially independent of the core CCR4–NOT complex.

Together, our findings show that the CCR4–NOT complex has a major role in regulating the transcriptome of human cells. The results demonstrate that the functional contributions of individual subunits are shaped by the structural organization of the complex. While depletion of the CNOT1 scaffold and CNOT7&8 deadenylases produced the most extensive transcriptomic effects, each module contributed unique regulatory signatures, revealing both cooperative and specialized functions within the CCR4–NOT complex.

### Identification of transcripts bound and regulated by CCR4–NOT

To identify RNAs associated with the CCR4–NOT complex, we used formaldehyde-crosslinked RNA immunoprecipitation RNA-seq (RIP-seq) to capture CCR4–NOT-bound transcripts in vivo (71). To do so, HCT116 cells were engineered to stably express a doxycycline-inducible V5-tagged CNOT7 catalytic mutant (D40A; V5-CNOT7^MUT^) (**Fig. 3A**). This mutant is incorporated into the CCR4–NOT complex but is unable to deadenylate target transcripts, thereby stabilizing associated RNAs and enhancing their recovery. Following crosslinking and anti-V5 immunoprecipitation, co-purifying RNAs were sequenced and mapped to the transcriptome. Successful incorporation of the V5-CNOT7^MUT^ into the CCR4–NOT complex was confirmed by co-immunoprecipitation of CNOT1 (**Fig. 3A**). Using a >1.5-fold enrichment relative to input RNA and Benjamini–Hochberg adjusted p-value of <0.01, we identified transcripts from 2, 930 genes enriched with the CCR4–NOT complex in HCT116 cells (**Fig. 3B**). While crosslinking approaches could conceivably link RNAs through a bridging protein, herein, we refer to CNOT7-bound mRNAs as those associated, indirectly and directly, with CNOT7.

**Figure 3.**
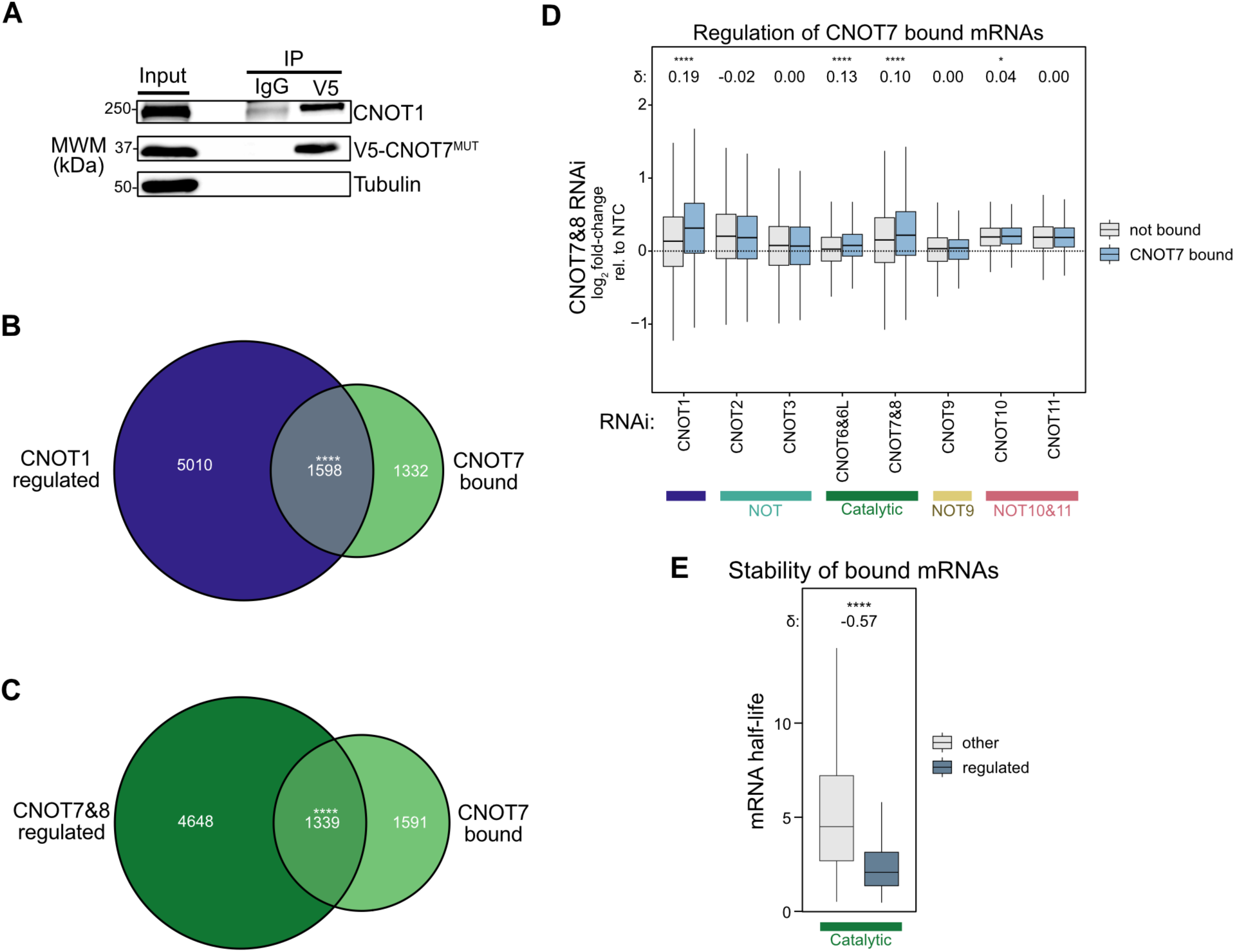
Identification of transcripts associated with CNOT7 and whose steady-state levels increased following CCR4–NOT depletion. (**A**) Western blot validation of V5-CNOT7^MUT^ immunoprecipitation from RIP-seq experiment; CNOT1 served as a positive control and tubulin served as a negative control. (**B-C**) Venn diagrams showing the overlap between transcripts bound by V5-CNOT7^MUT^ in RIP-seq (CNOT7 bound) and transcripts upregulated following (**B**) CNOT1 RNAi depletion (CNOT1 regulated) or (**C**) CNOT7&8 RNAi depletion (CNOT7&8 regulated). Each overlap was significant (p < 0.0001 = ****) based on Fisher’s exact test with p-values corrected via the Benjamini-Hochberg method. (**D**) Boxplots of the log_2_ fold-changes in transcript abundance in response to each CCR4–NOT subunit depletion in relation to transcripts binned as CNOT7-bound or not detectably bound. (**E**) Boxplots of mRNA half-life for transcripts regulated by the catalytic module (upregulated in CNOT1 and CNOT7&8, and associated with CNOT7), compared to all other transcripts. Pairwise comparisons were performed using a two-sided Wilcoxon rank-sum test with p-values adjusted using the Benjamini-Hochberg method. In all panels, p < 0.05 = *, p < 0.01 = **, p < 0.001 = ***, p < 0.0001 = ****. Cliff’s delta (δ) for each comparison noted.

To determine which of these bound RNAs are regulated by CCR4–NOT, we compared the RIP-seq dataset with transcripts whose abundance changed in response to depletion of CCR4–NOT subunits. Approximately half of the CNOT7-bound transcripts overlapped with genes upregulated upon depletion of either CNOT1 (54%, odds ratio 2.1) or CNOT7&8 (45%, odds ratio 1.6) (**Fig. 3B, C, Supplementary File S5**), indicating that a substantial portion of CCR4– NOT-bound RNAs are actively regulated by the complex.

To extend this analysis across all subunits, transcripts were binned according to CNOT7 binding status and compared with the distribution of expression changes observed in each knockdown. CNOT7-bound transcripts exhibited significantly greater upregulation following depletion of CNOT1, CNOT6&6L, and CNOT7&8 (**Fig. 3D**). For comparisons of distributions, the Cliff’s δ statistic was used to quantify how strongly the distributions differed. Cliff’s δ is a nonparametric effect-size statistic ranging from |δ| = 0 (indicating no difference) to |δ| = 1 (indicating no overlap) (75). We further considered the subset of transcripts that were upregulated by CNOT1, CNOT7&8, and associated with CNOT7. These high confidence targets have substantially shorter half-lives in normal wild-type conditions relative to all other transcripts (Cliff’s δ -0.57) (**Fig. 3E**). Together, these results identify a large set of RNAs that are both physically associated and functionally regulated by CCR4–NOT, consistent with its role in promoting their RNA deadenylation and decay.

### CCR4–NOT regulates networks of genes involved in proteostasis, stress responses, and survival

To gain insight into the biological pathways and processes regulated by CCR4–NOT, we performed GSEA to identify GO terms associated with up- and downregulated genes (**Fig. 4A**). Consistent with a high correlation between CCR4–NOT subunits, GO terms related to amino acid activation, inflammatory response, apoptotic signaling, and ER stress were upregulated following depletion of CNOT1, CNOT2, CNOT3, CNOT6&6L, and CNOT7&8. These same GO terms were not enriched with depletion of CNOT9, CNOT10, or CNOT11, and further terms such as intermediate filaments and tRNA aminoacylation were even downregulated following CNOT9 RNAi. Notably, GO terms related to DNA replication and chromosome segregation were enriched among genes downregulated following CNOT2 and CNOT3 depletion. This is likely due to the decrease in histone mRNA levels when these subunits were depleted (**Supplementary File S4, S5**). The same terms were enriched among upregulated genes following depletion of other subunits. While more specialized enriched processes included lipid modification (CNOT1, CNOT2, CNOT3, and CNOT7&8), and Rap signaling (CNOT10, CNOT11, CNOT6&6L, and CNOT7&8), GSEA reflects a general stress response following depletion of the catalytic and NOT modules.

**Figure 4.**
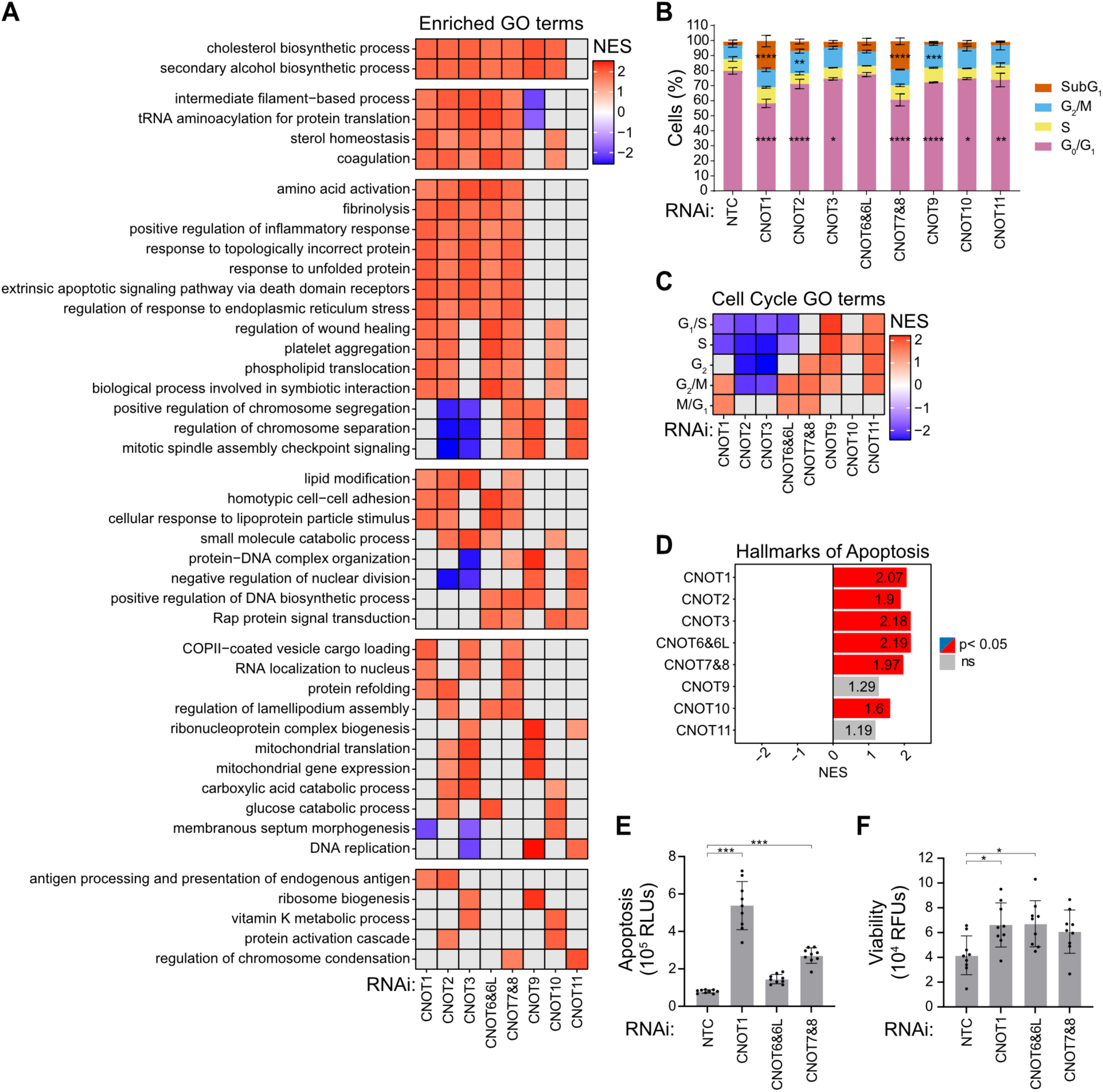
CCR4–NOT regulates networks of genes involved in stress responses, cell proliferation, and survival. (**A**) Heatmap showing top significantly enriched Gene Ontology (GO) Biological Process terms among upregulated genes (log₂ fold-change > 0; p < 0.05), for each CCR4–NOT subunit knockdown, as determined by Gene Set Enrichment Analysis (GSEA). Color reflects the Normalized Enrichment Score (NES) for significantly enriched terms. Grey denotes non-significant enrichment. Significance was calculated by permutation tests with Benjamini-Hochberg corrected p-values (p^adj^ <0.05). (**B**) Stacked bar graphs showing the percentage of cells at each stage of the cell cycle (G_0_/G_1_, S, G_2_/M, and SubG_1_) for each subunit knockdown condition, measured by propidium iodide staining and flow cytometry. n=3 biological replicates; +/- SD. For significance calling, p < 0.05 = *, p < 0.01 = **, p < 0.001 = ***, p < 0.0001 = **** based on ordinary one-way ANOVA and Tukey test for multiple comparisons. (**C**) Heatmap of NES from GSEA considering transcripts associated with the indicated cell cycle phase. (**D**) GSEA for the hallmarks of apoptosis gene sets. (**E**) Apoptosis and (**F**) cell viability measured following RNAi depletion of CNOT1, CNOT6&6L, CNOT7&8, or NTC. n=9 biological replicates; +/- SD. Significance thresholds as in panel C.

GSEA was complemented with GO enrichment analysis, which only considered the genes that were significantly upregulated following depletion of each CCR4–NOT subunit (**Fig. S4A-D, Supplementary File S5**). Consistent with GSEA, depletion of CNOT1 and CNOT7&8 affected broad GO categories, including macroautophagy, regulation of autophagy, and response to ER stress. In addition, terms related to Cajal bodies were highly enriched, due to upregulation of the TriC/CCT Complex. As a form of gene set analysis (76), over-representation analysis could be particularly informative for the CNOT6&6L and CNOT9 RNAi conditions that produced limited transcriptome-wide effects. Among genes upregulated following CNOT6&6L depletion, terms including microautophagy, nucleophagy, serine transport, and actomyosin contractile ring organization were enriched (**Fig. S4C**). Among genes upregulated following CNOT9 depletion, GO terms related to DNA replication and double-strand break repair were enriched (**Fig. S4D**).

As mitosis and DNA replication terms emerged as prominent features in GSEA and GO term analyses (**Fig. 4A, S4A-D**), we investigated how CCR4–NOT depletion influences cell cycle states. We performed flow cytometry analysis of DNA content following the depletion of each subunit (**Fig. 4B**). Depletion of CCR4–NOT subunits was confirmed by western blot (**Fig. S5A**). A reduction in the G_0_/G_1_ population was observed for all subunits except CNOT6&6L, accompanied by an increase in the SubG_1_ fraction for CNOT1 and CNOT7&8, and an increase in G_2_/M for CNOT2 and CNOT9 (**Fig. 4B**). No significant changes were detected in S phase under any condition. GSEA further revealed changes in cell cycle gene expression programs following depletion of specific CCR4–NOT subunits (**Fig. 4C**), which may be a consequence of altered cell cycle. Specifically, CNOT9, CNOT10, and CNOT11 showed positive enrichment for transcripts associated with G_1_/S and S phase, whereas all the remaining subunits exhibited negative enrichment for these sets. CNOT2 and CNOT3 were the only subunits that displayed significant downregulation of transcripts associated with G_2_ and G_2_/M, while the other subunits showed positive enrichment for these phases. All subunits showed positive enrichment for M/ G_1_, with CNOT1 and the catalytic modules passing the significance cutoff. Overall, these results indicate widespread alterations in cell cycle phase distributions following depletion of the individual CCR4-NOT subunits.

Because the proapoptotic signaling pathway was upregulated by CCR4–NOT depletion (**Fig 4A**), we next examined whether disruption of CCR4–NOT function induced the apoptosis pathway. We observed that depletion of CNOT1 and CNOT7&8 increased the proportion of cells exhibiting a SubG_1_ DNA content profile (**Fig. 4B**), a hallmark of apoptotic DNA fragmentation (77). Consistent with this finding, gene set enrichment analysis revealed upregulation of apoptosis-related hallmark genes following depletion of multiple CCR4–NOT subunits (**Fig. 4D**). We also detected increased caspase activity upon depletion of CNOT1 and, to a lesser extent, the catalytic subunits (**Fig. 4E**). Despite the induction of apoptosis-associated gene expression and caspase activity under these conditions, the overall cell viability remained largely unchanged (**Fig. 4F and Fig. S1A**), indicating that extensive cell death had not yet occurred.

Given the effects of CCR4–NOT depletion on cell cycle regulation and apoptosis, we next asked whether these perturbations extended to broader cancer-associated transcriptional programs.

We therefore performed GSEA for the hallmarks of cancer (**Fig. S5B**). The ten hallmarks of cancer were generally upregulated in response to knockdown of CCR4–NOT subunits. An exception is CNOT9 depletion, which caused downregulation in select cancer hallmarks, including sustained angiogenesis and proliferative signaling (**Fig. S5B**).

The observed GO terms related to mitochondrial gene expression (**Fig. 4A**) prompted us to measure potential changes in mitochondrial function. Using a luminescence-based assay that measures ATP production and cell membrane integrity in growth conditions that favor oxidative phosphorylation, we did not observe substantial changes in mitochondrial function in the CCR4– NOT subunit knockdowns (**Fig. S6A, S6B)**.

Together, these findings reveal that CCR4–NOT coordinates multiple interconnected biological programs that govern cellular homeostasis, including proteostasis, stress responses, cell cycle, and cell survival. While CNOT1 and the CNOT7&8 deadenylases exert broad regulatory effects consistent with their central roles in the complex, individual subunits contribute to specialized functions that can shape distinct gene expression and physiological outcomes.

### Distinct contributions of CCR4–NOT subunits to poly(A) length control

To examine the role of CCR4–NOT in deadenylation, we measured changes in poly(A) length caused by depletion of individual CCR4–NOT subunits. Nanopore direct RNA sequencing was performed on two biological replicates of RNA from the samples used for RNA-seq in **Figure 1**, and poly(A) lengths were estimated using the Dorado algorithm. Depletion of CNOT1 or CNOT7&8 deadenylases caused a shift in the bulk distribution of poly(A) tail lengths towards longer tails relative to NTC treated cells (**Fig. 5A**), and differences in poly(A) length distributions across all samples relative to NTC controls were assessed using Cliff’s δ **(Fig. S7A**). Median bulk poly(A) tail lengths increased from 76-83 nt in the NTC samples to 89-105 nt in the CNOT1 depleted samples and 105-112 nt for CNOT7&8-depleted samples. In comparison, depletion of CNOT6&6L deadenylases had little effect, exhibiting median bulk poly(A) length of 78-91 nt (**Fig. 5A, S7A**). These observations demonstrate the importance of CNOT7&8 in bulk deadenylation. Depletion of the other subunits had little to no effect on bulk length of poly(A) tails (**Fig. S7A**).

**Figure 5.**
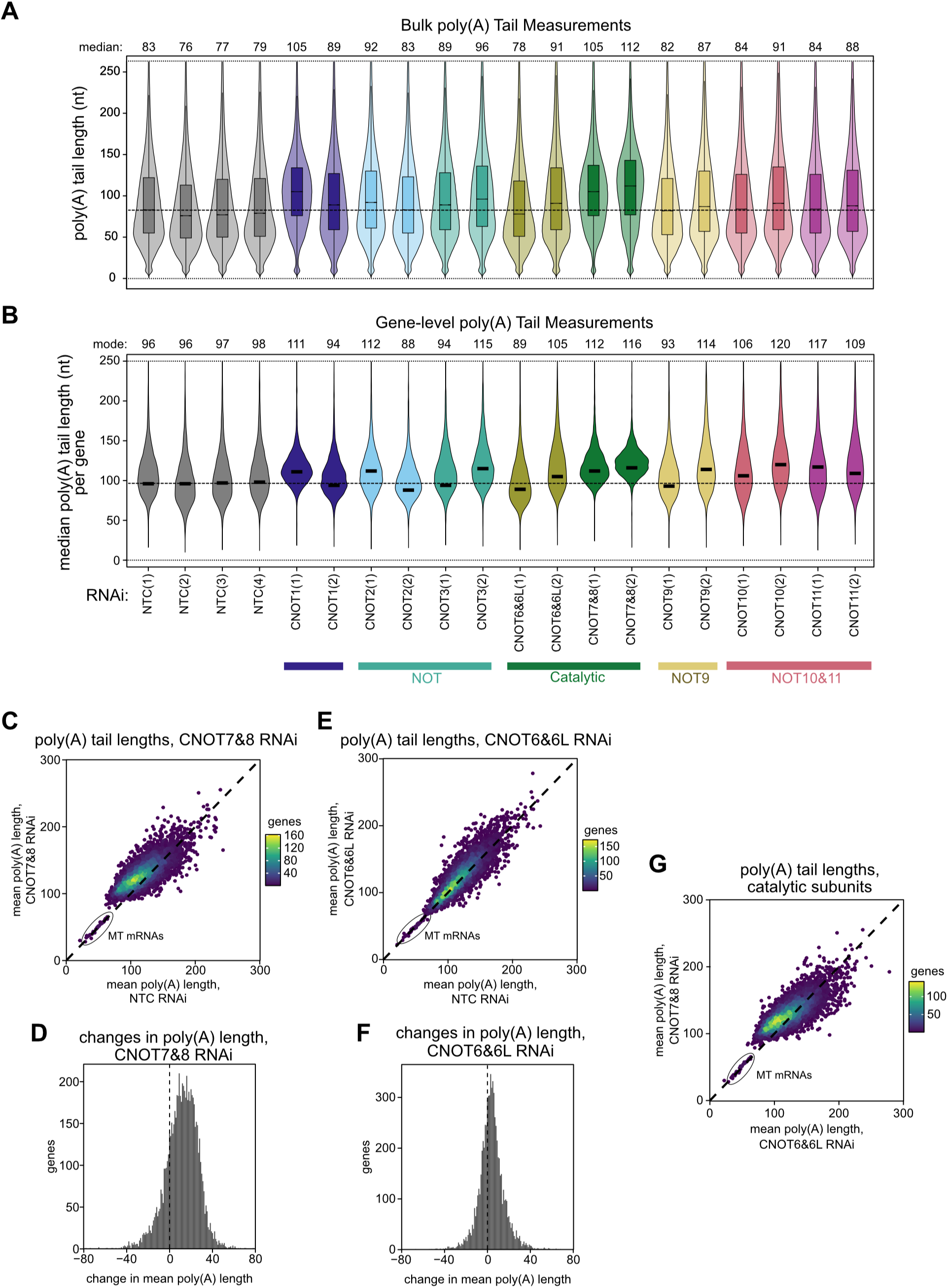
Distinct contributions of CCR4–NOT subunits to poly(A) length control. (**A**) Violin plots and boxplots showing the distributions of bulk poly(A) tail lengths when CCR4–NOT subunits are depleted compared to non-targeting control (NTC), as measured by nanopore sequencing. The dotted line marks the median value for the NTC replicate 1 for reference. (**B**) Violin plots showing the distributions of median poly(A) tail lengths for all profiled genes. Only genes with ≥10 measurements in all samples are considered. Black bar highlights the most often occurring median poly(A) tail length. Scatterplots of mean poly(A) tail length in NTC and CNOT7&8 RNAi samples (**C**), NTC and CNOT6&6L RNAi samples (**E**), and between catalytic subunits (**G**). Histograms of poly(A) tail length differences upon CNOT7&8 depletion (**D**) and CNOT6&6L depletion (**F**).

Measurements of bulk poly(A) lengths can be biased by highly expressed mRNAs. In our dataset, the five most highly profiled genes accounted for 5.5-7.8% total poly(A) measurements across the samples, with the most highly profiled genes (*ACTG1* or *EEF1A1*) accounting for 1.7- 3.0% alone (**Fig. S7B**), consistent with previous observations (78). Therefore, we assessed the distribution of median poly(A) lengths at the gene level (**Fig. 5B**, and Cliff’s δ reported in **Fig. S7C**). For accurate estimates, we considered only the 6, 951 genes profiled with 10 or more poly(A) measurements in all samples. As with bulk poly(A) measurements, CNOT7&8 depletion increased the median poly(A) tail of most genes. Compared to control samples, depletion of CNOT7&8 generally increased mean poly(A) tail lengths by approximately 20 nucleotides (**Fig. 5C-D**). A notable exception are mitochondrially-encoded mRNAs, with short mean tail lengths, that were insensitive to CNOT7&8 depletion; this is consistent with compartmentalization of these transcripts inside mitochondria (79). We also examined the relationship of poly(A) tail length to responsiveness of transcripts to CCR4–NOT. Depletion of CNOT7&8 altered the mRNA levels for transcripts that normally exhibit a broad range of mean poly(A) lengths, spanning ∼50-200 nucleotides (**Fig. S7D**). In comparison, depletion of CNOT6&6L slightly shifted mean poly(A) tail lengths (**Fig. 5E-F**), and to a lesser extent than depletion of CNOT7&8 (**Fig. 5G**). Finally, in addition to CNOT7&8 RNAi, depletion of CNOT10 or CNOT11 increased tail lengths at the gene level, whereas depletion of other CCR4–NOT subunits did not consistently alter tail length (**Fig. S7A, S7C**). These results demonstrate that CCR4–NOT deadenylates a broad set of transcripts and support the conclusion that CNOT7&8 are the predominant deadenylases in the complex.

### CCR4–NOT subunits differentially contribute to translation-dependent mRNA decay

Because CCR4–NOT function is linked to translation-dependent mRNA decay mechanisms, we examined whether features related to translation were associated with CCR4–NOT mediated regulation. To assess relationships to translation efficiency, we used previously reported measurements that were made by ribosome profiling in normal wild-type HCT116 cells (72). We also examined the relationship of codon optimality (80, 81) by calculating the fraction of optimal codons in each coding sequence using the codon stability coefficient (CSC), an experimentally derived metric that relates codon usage to mRNA stability (73). Most human genes have between 25-75% optimal codons. We note that translation efficiency and codon optimality parameters are only weakly correlated across all profiled genes (R^2^ 0.03, p-value < 0.001).

We investigated whether transcripts regulated by individual subunits and modules of CCR4– NOT subunits differed in these metrics. Transcripts regulated by the NOT module (upregulated in response to CNOT1, CNOT2, and CNOT3 RNAi) have significantly higher translation efficiency under normal conditions relative to all other genes. This difference is classified as small based on Cliff’s δ of 0.20 (**Fig. 6A**). Consistent with this, CNOT2 upregulated genes (δ 0.17) and CNOT3 upregulated genes (δ 0.23) have higher translation efficiencies (**Fig. 6B**).

**Figure 6.**
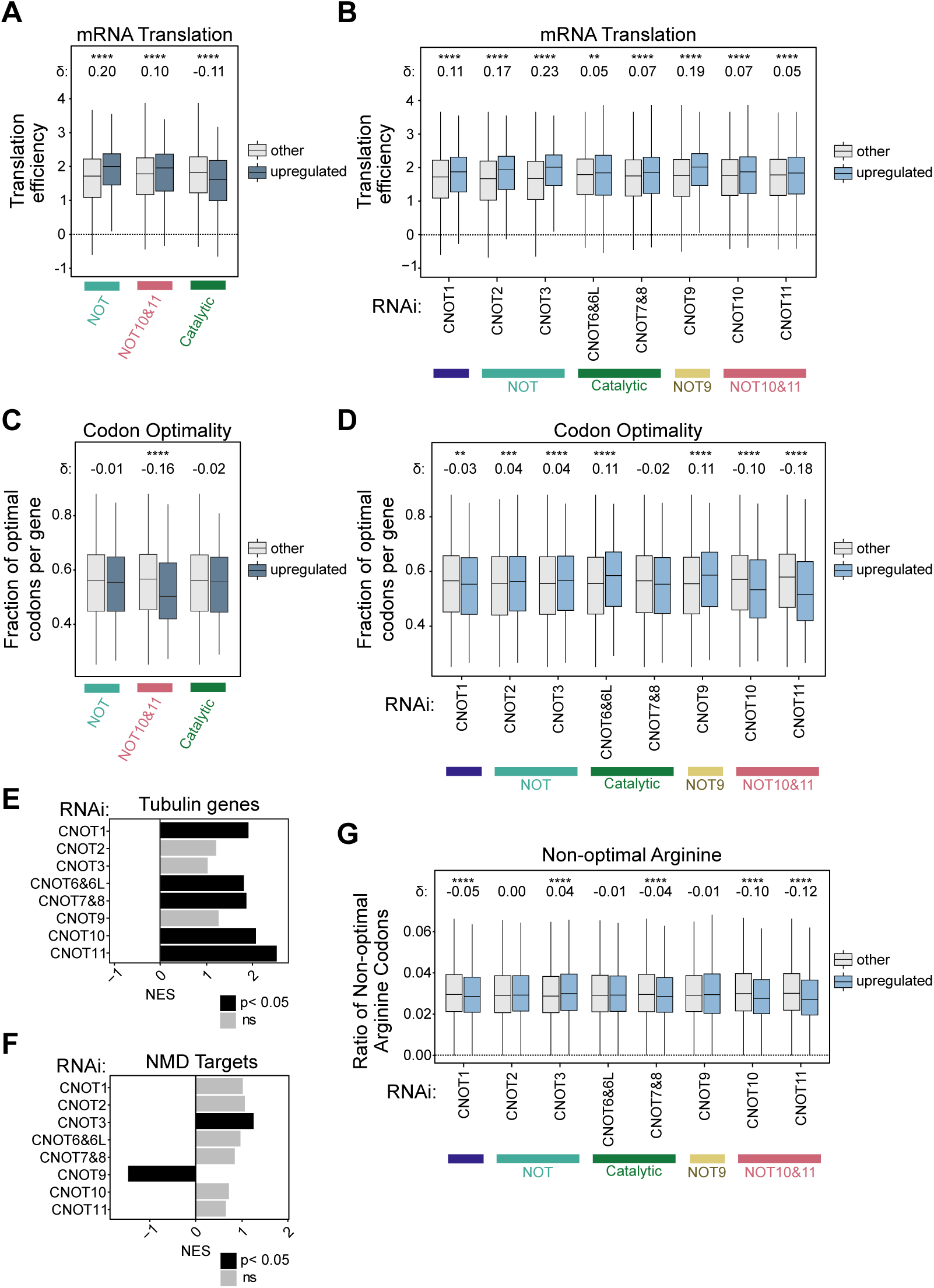
CCR4–NOT subunits differentially contribute to translation-dependent mRNA decay. Boxplots showing translation efficiency (**A-B**) and fraction of optimal codons (**C-D**), for transcripts binned by co-regulation by module subunits (**A, C**) or individual subunit knockdowns (**B, D**). In these panels, co-regulated transcripts (dark blue) and all upregulated transcripts (blue) were compared to all other respective genes (grey). GSEA for (**E**) tubulin genes and (**F**) Nonsense-mediated mRNA decay targeted gene sets (NMD) for each RNAi knockdown condition. (**G**) Fraction of non-optimal arginine codons for transcripts that are upregulated (blue) or not significantly upregulated (grey) in response to knockdown of each respective CCR4–NOT subunit. The significance of all comparisons between distributions was determined via two-sided Wilcoxon rank-sum tests with p-value adjusted via the Benjamini–Hochberg method. In all panels, p < 0.05 = *, p < 0.01 = **, p < 0.001 = ***, p < 0.0001 = ****.

Genes upregulated following the depletion of CNOT9 also have higher translation efficiency in normal conditions by a small degree (δ 0.19) (**Fig. 6B**). The relationship of translation efficiency relative to responsiveness to depletion of the other subunits were deemed negligible based on the Cliff’s δ <0.147. Overall, these comparisons indicate that the NOT module and CNOT9 may preferentially regulate stability of mRNAs that tend to be more efficiently translated.

Transcripts affected by the depletion of the NOT10–11 module (upregulated following CNOT10, CNOT11, and CNOT1 RNAi) exhibit a smaller fraction of optimal codons (δ -0.15) than all other transcripts (**Fig. 6C**). A similar relationship was observed for the individual CNOT11 depletion (**Fig. 6D**). Differences for the remaining subunits were negligible, including the catalytic module. These results suggest the NOT10–11 module may participate in selective targeting of mRNAs with suboptimal codons.

One mechanism of translation-dependent recruitment mechanism involves recognition of nascent amino-terminal tubulin peptides by the factors TTC5 and SCAPER, which recruit CCR4–NOT through interactions with CNOT10 and CNOT11 (55). GSEA of tubulin genes revealed upregulation of tubulin transcripts across multiple knockdowns, with the strongest effects observed following depletion of CNOT10, CNOT11, CNOT1, CNOT6&6L, and CNOT7&8 (**Fig. 6E**). Of the five classes of tubulins, transcripts from four out of five alpha and all eight beta tubulin genes increased in response to CCR4–NOT depletion (**Fig. S8**). In contrast, the delta, gamma, and epsilon tubulin genes exhibited little or no response to CNOT10 and CNOT11 depletion, consistent with their lack of the amino-terminal peptide sequence that recruits TTC5 and SCAPER. These findings strengthen the prevailing model of nascent peptide-mediated mRNA decay in tubulin homeostasis and provide a comprehensive assessment of the contribution of CCR4–NOT subunits to that regulation.

Translation-dependent deadenylation is thought to contribute to nonsense-mediated decay (NMD). To examine the involvement of CCR4–NOT in this pathway, we performed GSEA using a curated set of endogenous NMD targets (61). NMD substrates were significantly enriched among transcripts that increased following CNOT3 depletion (**Fig. 6F**), suggesting that CNOT3 contributes to degradation of these targets. In contrast, depletion of CNOT9 produced the opposite trend, raising the possibility that CNOT9 exerts a distinct, potentially antagonistic, role in NMD that warrants future investigation. Although transcripts upregulated following depletion of the remaining CCR4–NOT subunits also showed positive enrichment for NMD targets, these effects did not reach statistical significance. Overall, these findings support a role for CCR4– NOT in decay of transcripts targeted by NMD.

One additional mechanism of translation-dependent mRNA decay, P-site tRNA-mediated decay, involves recruitment of CCR4–NOT to ribosomes decoding non-optimal codons (CGG, CGA, and AGG) (54). We searched for signatures of this pathway in our dataset. These non-optimal arginine codons occur infrequently, with ratios between 0.02 – 0.04 for most genes (**Fig. 6G**), but are overrepresented in nuclear-encoded genes involved in mitochondrial translation (54). Though significant differences were detected between genes upregulated by CNOT1, CNOT3, CNOT7&8, CNOT10, an CNOT11 relative to all other genes, the effects were negligible. Thus, the impact of P-site tRNA-mediated decay was not evident in our data.

Together, these analyses provide a comprehensive view of how distinct CCR4–NOT subunits contribute to multiple forms of translation-dependent mRNA decay, including codon-mediated decay, nascent peptide-mediated decay, and nonsense-mediated decay.

### Identification of candidate recruitment factors linked to CCR4–NOT function

Multiple cis-acting RNA sequence motifs, and the trans-acting RNA-binding proteins (RBPs) that recognize them, recruit CCR4–NOT to selectively deadenylate target mRNAs (21). Using our functional datasets, we sought to identify additional recruitment factors that direct CCR4–NOT-mediated mRNA decay. To accomplish this, we leveraged a collection of human RBP:RNA interactions identified by crosslinking immunoprecipitation coupled with RNA sequencing (CLIP-seq) (62, 63). For 53 RBPs, we compared the bound RNA targets with transcripts that increased in abundance following depletion of individual CCR4–NOT subunits. From this analysis we identified RBPs whose target mRNAs were significantly enriched among transcripts upregulated upon CCR4–NOT subunit knockdown, suggesting that these RBPs may promote CCR4–NOT dependent degradation of their bound transcripts (**Fig. 7A, S9**).

**Figure 7.**
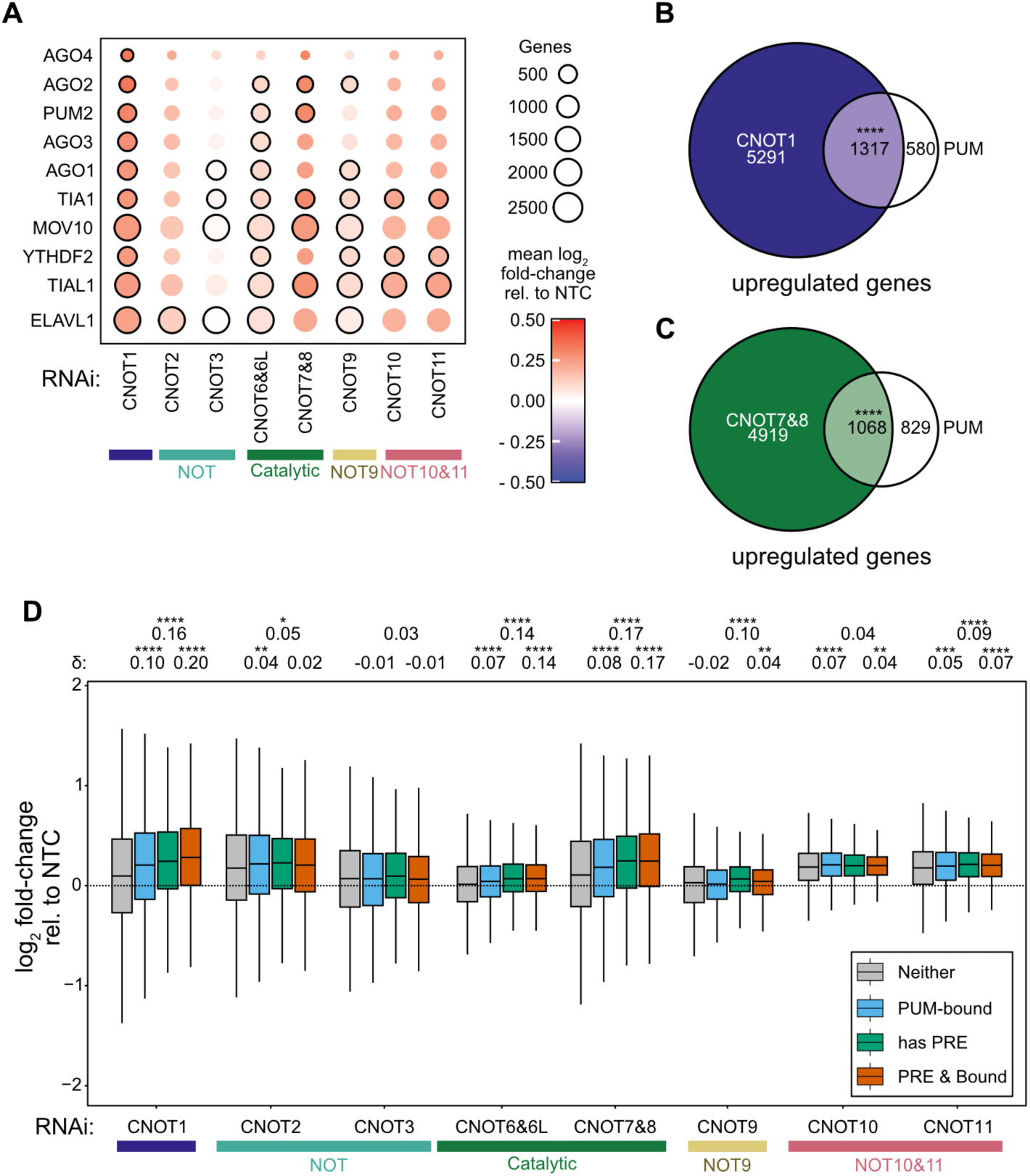
Identification of candidate recruitment factors linked to CCR4–NOT function. (**A**) mRNA targets of select RNA-binding proteins (RBPs) previously linked to CCR4–NOT regulation, identified using CLIP-seq data for HEK293T cells, were analyzed for differential expression in response to CCR4–NOT subunit knockdowns. The heat map color indicates mean log_2_ fold-change of the RBP targets in response to knockdown of the indicated subunit, circle size reflects the number of targets per RBP, black outline denotes statistical significance (p^adj^ <0.05). Significance was determined by a two-sided Wilcoxon test comparing target log_2_ fold-changes to all genes that are not bound by that RBP, and p-values corrected by the Benjamini-Hochberg method. The Venn diagram of mRNAs upregulated when PUM1&2 are depleted compared with those in the (**B**) CNOT1 or (**C**) CNOT7&8 depletion. Each overlap was significant (p < 0.0001 = ****) based on Fisher’s exact tests with p-values corrected via the Benjamini-Hochberg method. (**D**) Boxplots showing log_2_ fold-changes in RNA levels following depletion of each CCR4–NOT subunit. Genes were binned by PUM-bound, PRE-containing, or both PUM-bound and PRE-containing, and significance of differences determined via a two-sided Wilcoxon rank-sum tests with p-values adjusted via the Benjamini–Hochberg method (not significant [p>0.05] = ns, p < 0.05 = *, p < 0.01 = **, p < 0.001 = ***, p < 0.0001 = ****).

Several RBPs with established links to CCR4–NOT were identified in the results (**Fig. 7A**), including PUM2 (22, 39, 82), microRNA RISC (AGO1/2/3/4) (43–45, 83), and YTHDF2 (84, 85), providing validation of the approach. We also identified TIA1, TIAL1, and ELAVL1, which bind to destabilizing AU-rich RNA elements (86–90) (**Fig. 7A**). In addition, many RBPs with no previously recognized functional connection to CCR4–NOT were enriched in the transcripts upregulated by depletion of CCR4–NOT subunits, such as MOV10 (**Fig. 7A, S9**). For a subset of these RBPs, physical interactions with CCR4–NOT have been previously detected (24, 64) (**Fig. S9**). These results reveal potential new pathways for recruitment of the deadenylase complex.

The responses of transcripts bound by these RBPs exhibited subunit-specific patterns that mirrored the modular organization of the CCR4–NOT complex (**Fig. 7A, S9**). For example, PUM2 targets were preferentially upregulated following depletion of CNOT1, CNOT6&6L, and CNOT7&8 (**Fig. 7A**), consistent with previous reporter-based analyses (22, 82). Likewise, microRNA RISC targets responded to depletion of CNOT1, CNOT9, CNOT6&6L, and CNOT7&8 (**Fig. 7A**), in agreement with the established mechanism of microRNA-mediated mRNA decay (21). We also uncovered previously unrecognized subunit-specific trends.

Although YTHDF2 is known to interact with CNOT1 (84) and responded to its depletion (**Fig. 7A**), YTHDF2-bound transcripts were additionally upregulated by depletion of CNOT6&6L, CNOT9, CNOT10, and CNOT11, suggesting that these subunits also contribute to YTHDF2-directed CCR4–NOT-mediated mRNA decay.

Prompted by the enrichment of PUM2 targets among transcripts stabilized by CCR4–NOT depletion, we further investigated the functional relationship between Pumilio-mediated regulation and CCR4–NOT. We assessed the effect of depleting PUM1 and its paralog PUM2, as these proteins recognize the same RNA sequence motif, the Pumilio Response Element (PRE: 5′-UGUANAUA), and regulate overlapping sets of mRNAs. Consistent with a shared role in mRNA repression, transcripts upregulated following simultaneous depletion PUM1&2 (**Fig. S10A-C**) substantially overlap with those that are upregulated in response to CNOT1 depletion (**Fig. 7B**, odds ratio 3.7) or CNOT7&8 (**Fig. 7C**, odds ratio 2.3).

To focus on direct PUM1&2 targets, we integrated published datasets of PUM1/2-bound transcripts with mRNAs containing one or more PREs within their 3′ UTRs (22, 91–94). More than 1, 000 of these putative PUM1&2 targets were upregulated following depletion of CNOT1, CNOT2, and CNOT7&8, whereas fewer were affected by depletion of other CCR4–NOT subunits (**Fig. S10D**). Moreover, transcripts that contain PREs and that are also bound by PUM1/2 were more upregulated by depletion of CNOT1 and CNOT7&8 than transcripts without these features (**Fig. 7D**). To define a stringent set of shared direct targets, we intersected PRE-containing, PUM1/2-bound transcripts that were upregulated by PUM1&2 depletion with the mRNAs captured by CNOT7 RIP-Seq. This analysis identified 232 genes encoding proteins involved in signaling, transcription, membrane trafficking, and developmental pathways as high confidence targets of PUM1&2 that undergo CCR4–NOT-mediated repression. Together, these findings support a broad role for PUM1 and PUM2 as recruitment factors that direct CCR4– NOT-mediated degradation of target mRNAs (**Supplementary File S9**).

### Identification of new sequence motifs associated with subunit-specific CCR4–NOT function

We sought to identify cis-regulatory elements that confer responsiveness to CCR4–NOT. We performed de novo motif discovery using the FIRE algorithm, which applied stringent criteria to identify sequence motifs that were significantly overrepresented among transcripts that changed in abundance following depletion of individual CCR4–NOT subunits. Because determinants of deadenylation are typically located in 3′UTRs, our analysis focused on this region. This approach identified several uracil-rich motifs that were enriched among transcripts regulated by CNOT1, CNOT10, and CNOT11 **(Fig 8A)**. Similar motifs were enriched when different bin widths were used (data not shown).

**Figure 8.**
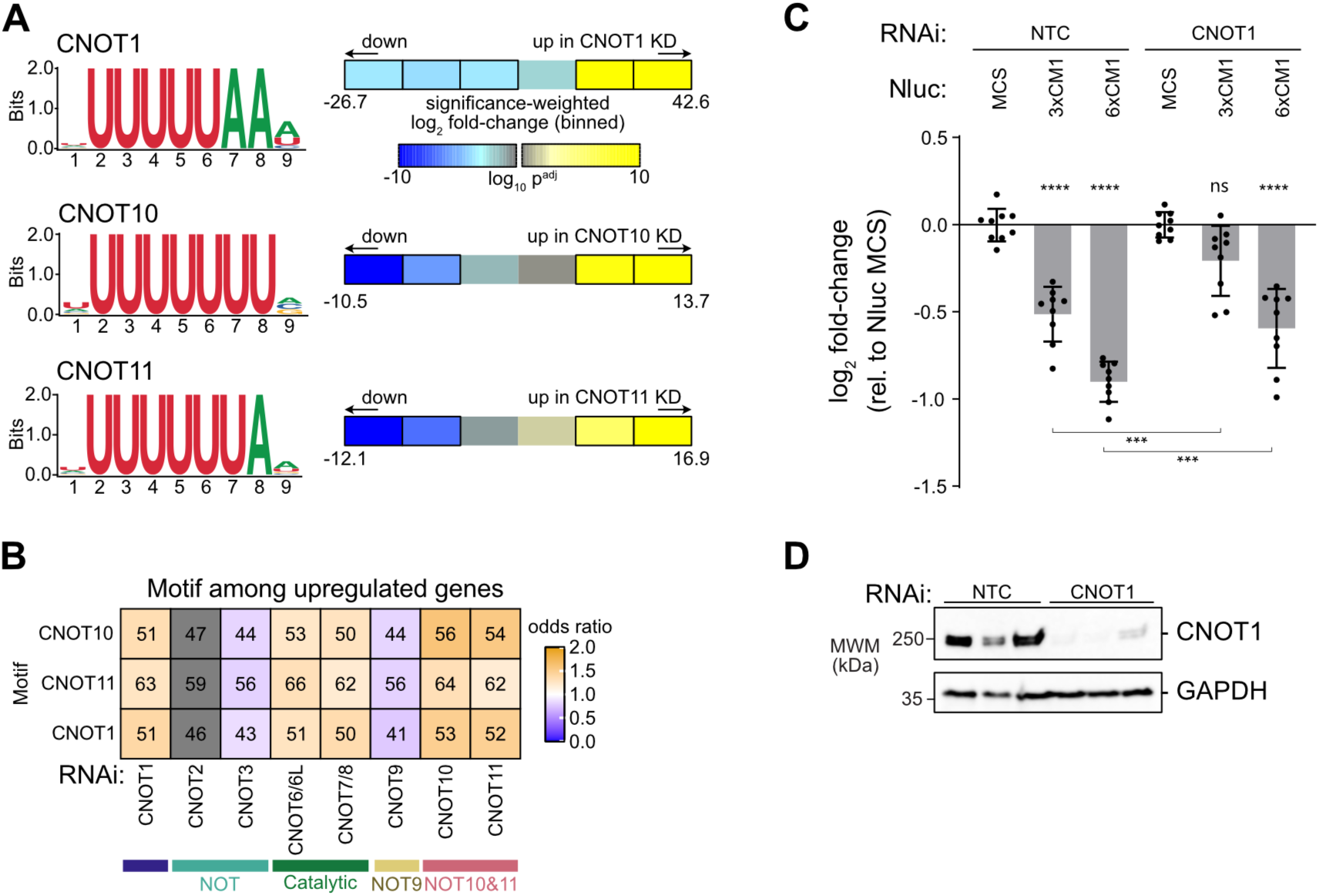
Identification of new sequence motifs associated with subunit-specific CCR4– NOT function. (**A**) FIRE output showing motif enrichment across six ordinal bins of significance-weighted log_2_ fold-changes. The motifs, presented as a sequence logo, are significantly over-represented in the bordered bins. The p-values were adjusted for multiple testing via the Bonferroni method. (**B**) The motifs identified by FIRE were profiled across upregulated genes. Numbers indicate the percentage of upregulated genes, following subunit knockdown, that contain at least one copy of the motif. The color scale reflects the odds ratio. Enrichment or depletion was determined using Fisher’s exact test and comparisons that did not pass the Benjamini-Hochberg adjusted p-value are masked in grey. (**C**) Repressive activity of the CCR4–NOT complex measured in HCT116 cells using Nano-luciferase reporters containing three or six motifs (Nluc 3xCM1 and 6xCM1, respectively) identified by FIRE within a minimal 3′UTR, relative to a control reporter containing a 3′UTR multiple cloning site (MCS). Reporter activity was quantified under non-targeting control (NTC) conditions and following RNAi knockdown of CNOT1. Firefly luciferase (Fluc) served as an internal control to assess variability in transfection efficiency. n=9; 3 experiments, each with 3 biological replicates; +/- standard deviation (SD). Significant differences determined by ordinary one-way ANOVA and Tukey test for multiple comparisons. (**D**) Verification of CNOT1 knockdown by western blot, with GAPDH as a loading control. For all panels: not significant [p>0.05] = ns, p < 0.05 = *, p < 0.01 = **, p < 0.001 = ***, p < 0.0001 = ****.

Because the U-rich motifs were highly similar, we examined their occurrence across the CNOT1, CNOT10, and CNOT11 regulated transcripts. A substantial fraction of these motifs occupied the same positions within 3′UTRs, indicating that FIRE had likely identified variants of the same underlying sequence element **(Fig. S11A)**. Consistent with this interpretation, the motifs were significantly enriched among transcripts upregulated following depletion of CCR4– NOT subunits with the exceptions of the NOT module and CNOT9 (**Fig. 8B**). Notably, a large percentage of upregulated transcripts (∼41-66%) contained one or more of these motifs (**Fig. 8B**).

To determine whether these U-rich motifs are functional, we defined a minimal consensus sequence, designated CNOT Motif 1 (CM1; 5′-UUUUUUUAAAA), that captures the shared features of the motifs associated with CNOT1, CNOT10, and CNOT11 regulation. We identified a CM1 site within the ETS1 3′UTR (Gene ID: 2113) and used its native sequence context to generate nanoluciferase reporters containing either three or six copies of the motif (Nluc 3xCM1 and Nluc 6xCM1). When expressed in HCT116 cells, both reporters exhibited significantly lower expression than the parental reporter containing only a multiple cloning site in the 3′UTR (**Fig. 8C**). Reporter activity was normalized to a co-transfected firefly luciferase control to account for differences in transfection efficiency.

Comparison of CM1 with previously characterized motifs revealed similarity to the cytoplasmic polyadenylation element (CPE; 5′-UUUUA_1-2_U), which is recognized by CPEB-family proteins (95). Among the CPEB family members, CPEB4 was the most highly expressed in HCT116cells based on our RNA-seq data. In addition, published PAR-CLIP data showed that CPEB4 binds the CM1-containing region of the ETS1 3′UTR (96), raising the possibility that CPEB4 mediates CM1-dependent repression. To test this hypothesis, we depleted either CNOT1 or CPEB4 and measured reporter activity. Knockdown of CNOT1 significantly reduced CM1-mediated repression (**Fig. 8C**), whereas depletion of CPEB4 had no detectable effect (**Fig.**

**S11B**). Efficient depletion of CNOT1 was confirmed by western blotting (**Fig. 8D**) and depletion of CPEB4 confirmed by RT-qPCR (**Fig. S11C**). Together, these results demonstrate that CM1 functions as a CNOT1-dependent regulatory element and suggest that it is recognized by one or more RNA-binding proteins distinct from CPEB4.

In summary, we identified a conserved U-rich motif that is enriched among transcripts regulated by CCR4–NOT and likely represents a common determinant of target specificity. This motif is sufficient to confer CNOT1-dependent repression, suggesting that it recruits CCR4–NOT through one or more unidentified RNA-binding factors. Importantly, the results demonstrate the usefulness of this dataset in identifying new CCR4–NOT regulatory mechanisms.

## Discussion

This study provides a comprehensive view of how individual CCR4–NOT subunits regulate gene expression in human cells. Depletion of every subunit altered numerous transcripts, underscoring the complex’s broad role in mRNA regulation. Subunits within the same biochemical module, including the NOT module, catalytic module, and the NOT10–11 module, affected largely overlapping transcript sets, consistent with their physical interactions and interdependent protein stability. Assessing all subunits highlighted the broad impact of the catalytic module and central CNOT1 scaffold, whereas other modules appear to have more specialized functions. Together, these findings demonstrate that the functional impact of CCR4– NOT reflects its modular biochemical architecture.

Among the subunits examined, depletion of CNOT1 or CNOT7&8 had the largest effects on transcript abundance, consistent with their central scaffolding and catalytic roles (22–24). As the CCR4–NOT complex can contain either CNOT6 or CNOT6L (CNOT6/6L) and CNOT7 or CNOT8 (CNOT7/8), the potential impact of the four possible combinations of enzymes remains unknown. Our data show that the combined CNOT7&8 depletion had a greater impact on transcript levels and poly(A) lengths than CNOT6&6L depletion, and transcripts upregulated after CNOT6&6L depletion largely overlapped with those affected by CNOT7&8. This agrees with prior work in mouse embryonic fibroblasts, where loss of CNOT7&8 increased transcript levels and poly(A) tail lengths to a greater extent than CNOT6&6L (97). These findings support a predominant role for CNOT7 and CNOT8 in CCR4–NOT-mediated deadenylation, with CNOT6 and CNOT6L providing complementary activity for a subset of transcripts. This interpretation is supported by a depletion-and-rescue study, which showed that loss of CNOT7&8 activity caused poly(A) tail lengthening, whereas loss of CNOT6&6L activity had little effect (18). By measuring both transcript abundance and poly(A) tail distributions across subunit depletions, our study extends those observations and connects the predominant contribution of CNOT7&8 to deadenylation with the broad effect on mRNA levels. It must be noted, however, that the architecture of the catalytic module may also contribute to the apparent difference.

Because CNOT7/8 bridge CNOT6/6L to the complex, CNOT7&8 depletion impairs both their own catalytic function and CNOT6/6L incorporation.

While the catalytic module and CNOT1 had the largest transcriptomic effects, depletion of other modules showed more restricted changes in a higher proportion of unique transcripts. CNOT9 depletion had a more limited impact overall and affected a distinct group of mRNAs. CNOT10 and CNOT11 regulated more than a thousand transcripts that did not respond to depletion of other subunits. While CNOT11 and CNOT2 depletions individually affected the largest number of unique targets (723 and 623, respectively), it is important to consider the interdependent stabilities of their protein partners within the modules. CNOT11 depletion also reduced CNOT10 protein, whereas CNOT2 knockdown also reduced CNOT1 and CNOT3 protein levels. As a consequence, the observed upregulation of target transcripts is likely capturing the co-depletion of their respective partners.

GSEA and GO analysis revealed that CCR4–NOT coordinates gene expression programs including proteostasis, endoplasmic reticulum and cellular stress responses, autophagy, metabolism, and signaling. Disrupting the complex also altered cell-cycle gene-expression programs and, particularly for CNOT1 or CNOT7&8 depletion, activated apoptotic responses. These findings are consistent with previous analyses of certain subunits that linked CCR4–NOT to apoptosis (97–102). Our study adds a systematic analysis of the contribution of all subunits in initiating the gene-expression changes associated with apoptosis. We also detected a general increase in expression of genes in each of the hallmarks of cancer upon depletion of CCR4– NOT subunits, except for CNOT9 regulated genes, which decreased in half of those categories. Gene expression of some processes, such as DNA replication or chromosome segregation, responded to a subset of depletions, further supporting module-specific functions.

Our findings implicate CCR4–NOT in mechanistically distinct forms of translation-coupled regulation (9, 103, 104). Previous studies linked CCR4–NOT to codon-mediated mRNA decay (51, 105). Depletion of CNOT1 was reported to preferentially stabilize transcripts enriched with non-optimal codons (24). CNOT3 is thought to sense ribosome state during decoding of non-optimal codons by binding to the ribosomal E-site when the A-site is empty (52, 53). Similarly, in P-site tRNA-mediated decay, CNOT3 can recognize the ribosomal E-site during decoding of specific non-optimal arginine codons (54). In our data, the relationships of non-optimal codons to changes in transcript levels were negligible for most of the CCR4–NOT subunits, with the exception of CNOT10 and CNOT11. We found that mRNAs upregulated by depletion of the NOT10–11 module have significantly fewer optimal codons. A potential role for CNOT10 and CNOT11 in sensing non-optimal codons is supported by their reported crosslinking to ribosomes decoding non-optimal codons (53). Together, the functional and physical connections of the NOT10–11 module to codon-mediated decay warrants future mechanistic investigation.

Our results also strengthen the connection of CCR4–NOT to the nonsense mediated decay pathway. NMD targets were most strongly stabilized after CNOT3 depletion, supporting a role for the complex in promoting their decay. CNOT9 depletion had the opposite effect, suggesting that CNOT9 may antagonize NMD through an unknown mechanism. The modest overall association between CCR4–NOT depletion and NMD substrates is consistent with mechanistic redundancy: in addition to deadenylation and exonucleolytic decay (49, 106–108), NMD can proceed through SMG5- and SMG6-mediated endonucleolytic cleavage (109–113), bypassing CCR4–NOT-mediated deadenylation.

CCR4–NOT also promotes tubulin mRNA autoregulation, a translation-dependent mechanism that maintains tubulin homeostasis (114). Nascent amino-terminal MREI/C peptides of alpha and beta tubulins are recognized by TTC5 and SCAPER, which recruit CCR4–NOT through the NOT10–11 module (55, 115–117). Consistent with that model, CNOT10 or CNOT11 depletion stabilized alpha and beta tubulin mRNAs, while similar effects following CNOT1, CNOT6&6L, or CNOT7&8 depletion demonstrate involvement of the broader complex. These effects occurred under normal growth conditions, indicating this pathway contributes to normal tubulin homeostasis and may respond more strongly during tubulin depolymerization (114). These transcriptome-wide data support nascent-peptide-directed CCR4–NOT recruitment and raise the possibility that analogous mechanisms respond to other nascent peptide signals.

Our results provide new insight into how CCR4–NOT selects its mRNA targets. We identified a U-rich element that defines a previously unrecognized route to CCR4–NOT-dependent repression. U-rich motifs were enriched among transcripts responding to CNOT1, CNOT10, and CNOT11 depletion, and the element was sufficient to confer CNOT1-responsive repression. The activity of this motif was not affected by depletion of CPEB4, an RBP with matching specificity, implicating an unidentified RBP or group of RBPs in its recognition and CCR4–NOT recruitment. Alternatively, the complex may engage this sequence directly, consistent with previous observations in both human and yeast systems showing that CCR4-NOT can bind to poly(U) RNA (35, 118).

The datasets reported here provide a versatile resource for studying CCR4–NOT-mediated regulation. The data can be used to interrogate whether a transcript, pathway, or RBP depends on CCR4–NOT and which subunits or modules contribute. Comparing RBP targets with transcripts affected by depletion of the corresponding interaction module can reveal the functional consequences of RBP:CCR4–NOT contacts. Furthermore, integration of our functional data with crosslinking and RNA coimmunoprecipitation (CLIP-seq) of CCR4–NOT bound transcripts can identify direct decay substrates (71). Our analysis of CLIP-seq datasets identified RBPs whose targets were enriched among CCR4–NOT-regulated mRNAs, including established recruitment factors such as PUM2, microRNA RISC, and YTHDF2, along with additional candidate RBPs. Many RBPs showed distinct relationships with individual CCR4– NOT subunits, providing transcriptome-wide evidence that recruitment factors can engage specific modules. Conversely, transcripts affected by all subunit depletions are strong candidates for regulation requiring the intact complex rather than specific components. Future integration with new RBP target datasets or established binding motifs can identify transcripts likely to be directly co-regulated by an RBP and CCR4–NOT. Likewise, identification of enriched motifs among co-regulated transcripts can reveal candidate cis-elements and RBPs when binding sites are unknown.

The utility of these datasets is illustrated by the known CCR4–NOT interaction with PUM1 and PUM2 (22, 82, 94). PUM-regulated, directly bound, PRE-containing transcripts strongly overlapped those regulated by CNOT1 and the catalytic module. These findings extend the mechanism of PUM-directed, CCR4–NOT-mediated repression that was established using reporter mRNAs and selected targets (22, 82) to a broader network enriched with targets encoding Ras/MAPK, small-GTPase, membrane-trafficking, and developmental signaling pathways.

This study has several limitations that also identify opportunities for future work. Analyses were performed in one human cell line, and the relative contributions of individual subunits may vary across cell types, developmental states, and physiological conditions. RNAi-mediated depletion is inherently incomplete and constrained by the time needed to achieve effective knockdown.

More rapid, extensive depletion could reveal additional regulatory effects, but it could also compromise viability and increase secondary consequences from disrupting this essential complex. Our analysis did not include CNOT4, as it is not a stable stoichiometric component of the human CCR4–NOT complex (17). Moreover, recent transcriptome-wide analysis indicates that CNOT4 has effects on mRNA metabolism distinct from those of the canonical CCR4–NOT complex (24). Our crosslinking RIP-seq approach examined only one of the four deadenylase subunits, CNOT7. Extending this approach to CNOT8, CNOT6, and CNOT6L could reveal additional direct targets. Our functional data was compared to a limited number of available RBP CLIP datasets conducted in different cell lines. CLIP approaches may be biased to highly expressed mRNAs, which are also more likely to pass the threshold of significance in differential gene expression studies. However, we note that differential gene expression was detected across expression levels, and significant overlap with CLIP datasets was observed for only specific knockdown conditions. Finally, our transcriptomic measurements captured steady-state RNA abundance and poly(A) length. Because deadenylation and decay are kinetic processes, approaches that directly measure transcript-specific rates should provide a more detailed view of how individual components determine mRNA fate. Together, kinetic measurements, broader direct-target identification, and analyses across diverse cellular contexts should further resolve the specialized and overlapping functions of CCR4–NOT components.

## Data Availability

The data underlying this article is available in the Gene Expression Omnibus, and can be accessed with accession number GSE320579 and GSE348528.

## Supplementary Data statement

Supplementary Data are available at *NAR* Online.

## Acknowledgements

We thank members of the Goldstrohm lab and Dr. Lydia Freddolino for insightful comments and suggestions throughout this study. Artificial intelligence was used for limited generation of and debugging of code, and for copy editing.

## Author Contributions Statement

K.M. conceptualization, investigation, formal analysis, resources, validation, conceptualization, visualization, writing, editing

M.N. data curation, formal analysis, investigation, methodology, resources, validation, visualization, writing, editing

R.C. data curation, visualization, formal analysis, validation, writing, editing

K.R. investigation, methodology, resources

Z.L. investigation, methodology, resources, formal analysis, validation

E.D. visualization, writing, analysis

M.F. funding acquisition, supervision, conceptualization, analysis, review

E.V. funding acquisition, supervision, conceptualization, analysis, review

J.B. funding acquisition, supervision, conceptualization, review, editing

A.C.G. Corresponding author, funding acquisition, project administration, supervision, conceptualization, visualization, writing, review, editing

