## Supplementary Figures for "Global Regulatory Functions of the Human CCR4–NOT Complex in mRNA Deadenylation and Decay"

**Supplementary Figure S1**

**McKenney et al.**

**A**

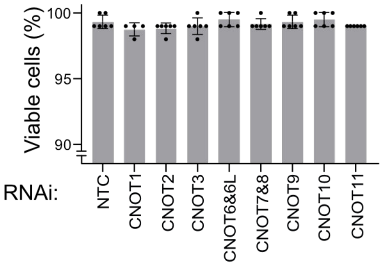


**B**

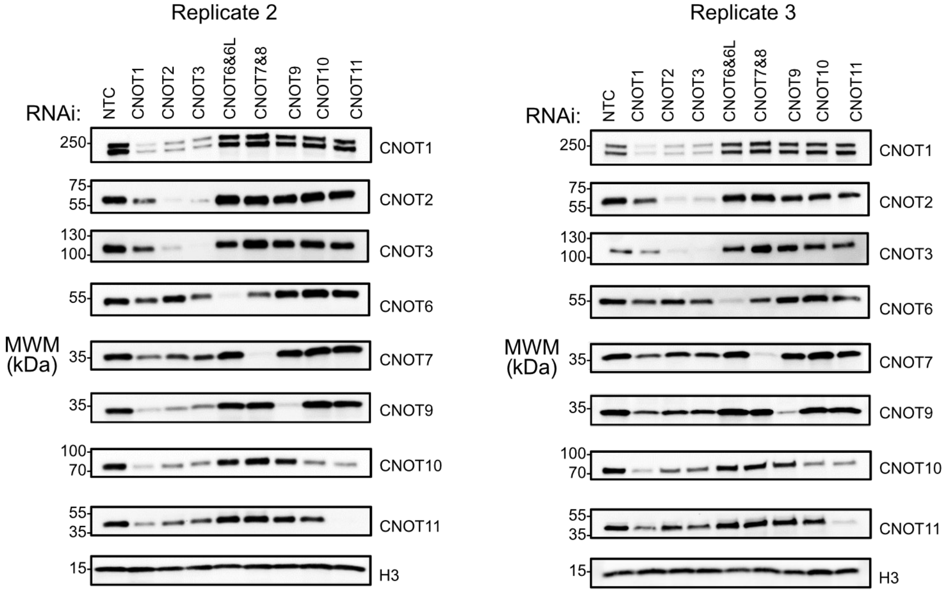

**Figure S1. Validation of CCR4–NOT subunit knockdown and assessment of cell viability.** (**A**) Cell viability following RNAi-mediated knockdown of each CCR4–NOT subunit and the non-targeting control (NTC) measured by trypan blue staining. Viability measurements were obtained from three biological replicates with two technical replicates each totaling six replicates per condition, except for CNOT1 which included four total replicates. Ordinary one-way ANOVA with Tukey's multiple-comparison test detected no significant differences in viability between the NTC and any CCR4–NOT knockdown condition. (**B**) Additional western blot replicates corresponding to Figure 1.

#### Supplementary Figure S2
#### McKenney et al.

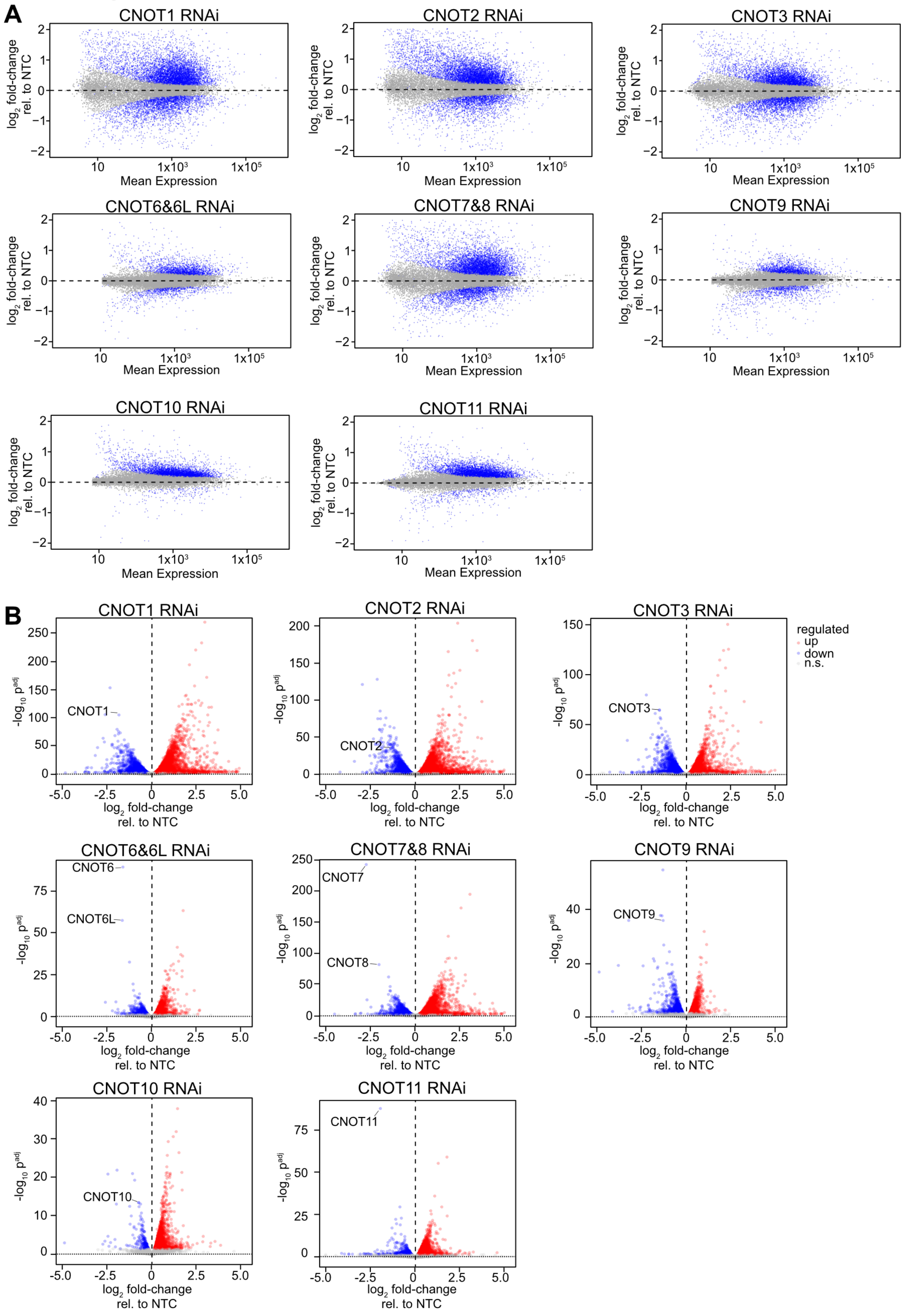

**Figure S2. Global gene expression changes following depletion of individual CCR4–NOT subunits.** (**A**) MA plots illustrating genome-wide expression changes for each CCR4–NOT subunit knockdown relative to the NTC, with $\log_2$ fold-change plotted as a function of mean expression level (DESeq2 baseMean). Genes exhibiting significant changes are shown in blue. (**B**) Volcano plots showing differential gene expression ($\log_2$ fold-change) versus significance (-$\log_{10}$ Benjamini–Hochberg-adjusted p-value ($p^{adj}$)) following depletion of each CCR4–NOT subunit, as indicated on each plot. Upregulated, downregulated, and nonsignificant genes are represented by red, blue, and gray points, respectively. Differential expression significance was defined as $p^{adj} < 0.05$.

**Supplementary Figure S3**

**McKenney et al.**

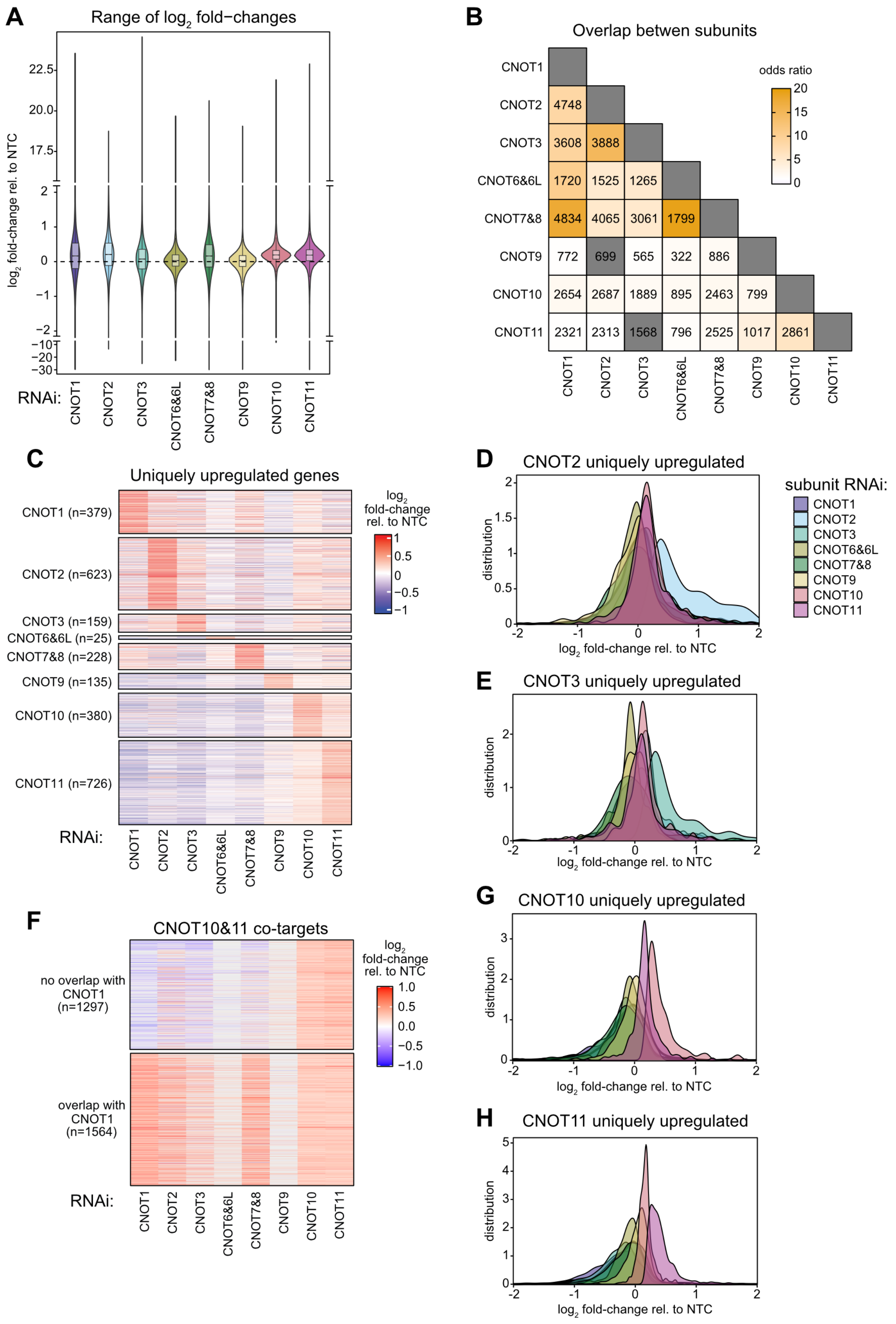

**Figure S3. Shared gene expression changes among CCR4–NOT subunits.** (**A**) Violin plots displaying the distribution of $log_2$ fold-changes for each CCR4–NOT subunit knockdown relative to the NTC. (**B**) Heat map showing the number of overlapping upregulated genes for each RNAi condition. Color intensity denotes odds ratio. Significance of overlap determined via Fisher's exact test with Benjamini-Hochberg method to correct for multiple testing. Overlaps that were not significant masked in grey. (**C**) Heat map showing the hierarchical clustering of $log_2$ fold-changes for transcripts uniquely upregulated in each subunit knockdown, compared with their expression patterns across the remaining subunit knockdowns. (**D-E**) Distributions of $log_2$ fold-change, relative to NTC RNAi, for depletion of each CCR4–NOT subunit. Distributions represented by kernel density estimate. Axis limited to -2 ≥ log2 fold-change ≥ 2. Genes considered were limited to those uniquely upregulated by CNOT2 (**D**) or CNOT3 (**E**). Heatmap of $log_2$ fold-changes of CNOT10 & CNOT11 co-targets that overlapped with CNOT1 upregulated genes (top) and genes that did not overlap (bottom). Non-significant changes masked with grey. (**G-E**) Distributions of $log_2$ fold-changes for genes uniquely upregulated by CNOT10 (**G**) or CNOT11 (**H**).

**Supplementary Figure S4**
**McKenney et al.**

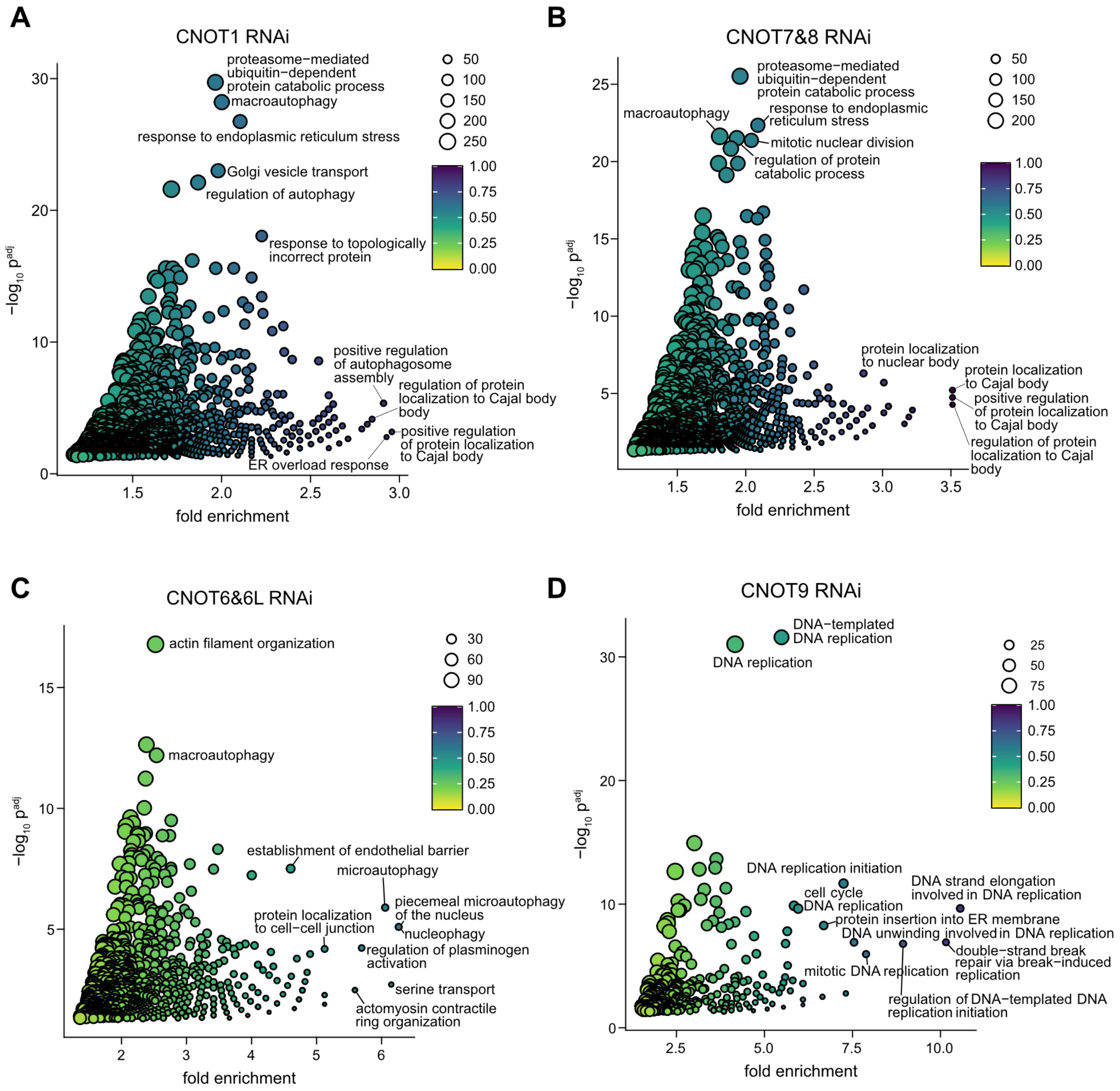

**Figure S4.** Scatterplots of fold enrichment versus significance ($-\log_{10} p^{adj}$) for GO terms that were significantly enriched among upregulated genes. Size denotes the number upregulated genes that are members of the GO term. Color denotes the fraction of GO term members that are also upregulated genes. Representative GO terms highlighted. Scatterplots were generated for CNOT1 (**A**), CNOT7&8 (**B**), CNOT6&6L (**C**), and CNOT9 (**D**) depletion. Full results of GO enrichment presented in **Supplementary File S5**.

**Supplementary Figure S5**

**McKenney et al.**

**A**

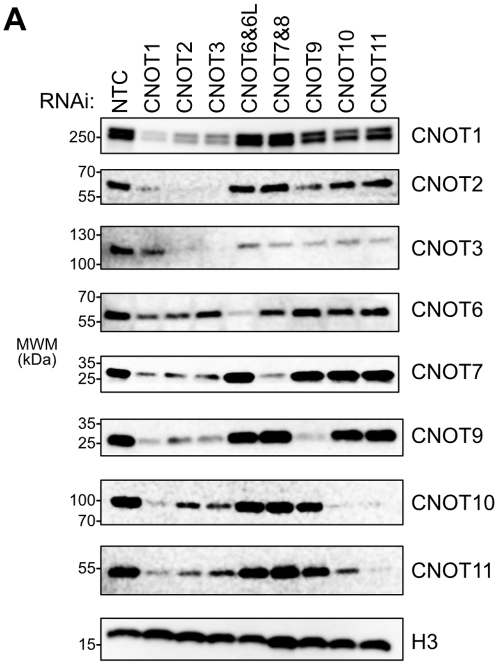


**B**

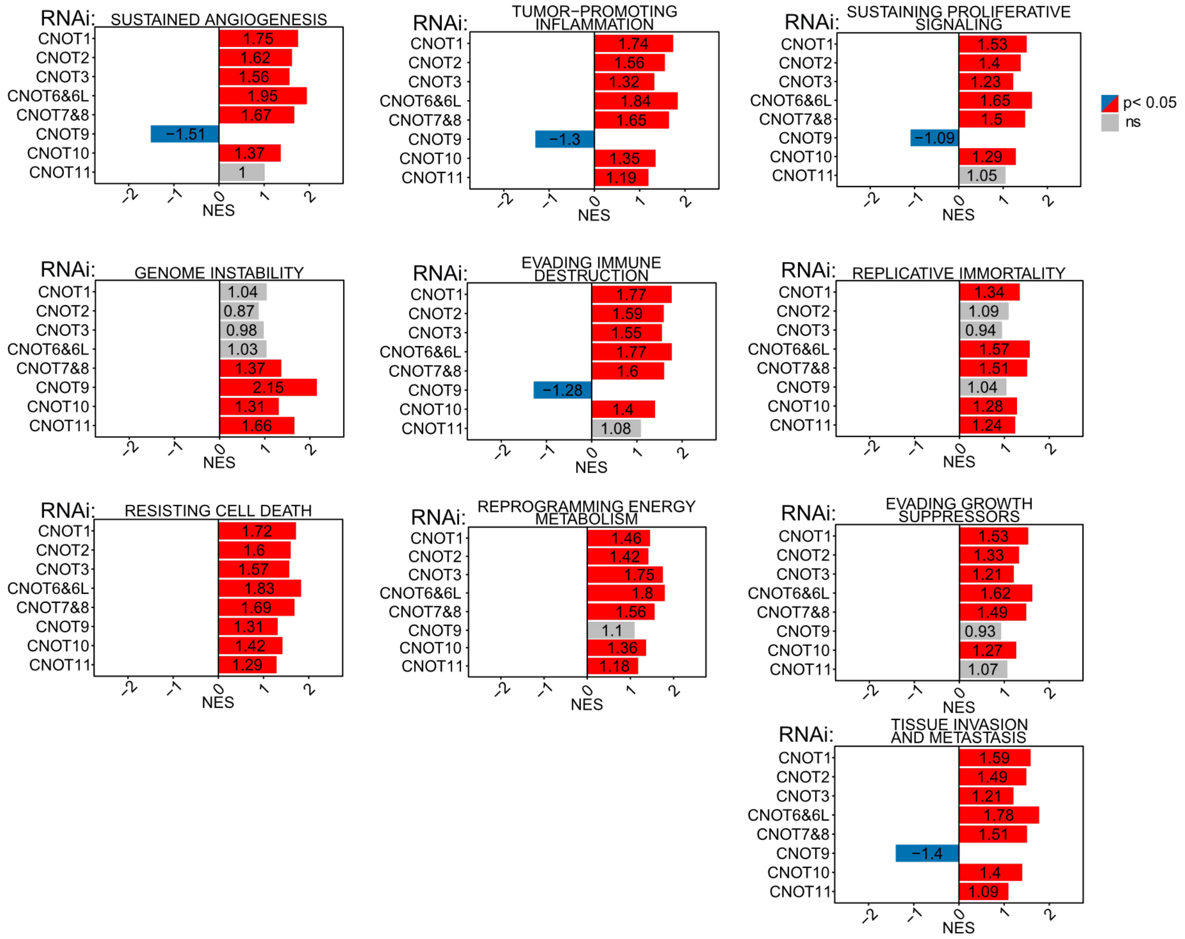

**Figure S5. Effects of CCR4–NOT subunit depletion on cancer-related gene signatures.** (**A**) Western blot confirmation of CCR4–NOT subunit RNAi knockdowns corresponding to the flow cytometry cell cycle analysis shown in **Figure 3B**. Histone 3 (H3) served as the loading control. (**B**) GSEA of transcripts involved in hallmarks of cancer. Enrichment significance was calculated by permutation test (two-sided), with multiple testing corrections using the Benjamini–Hochberg method.

**Supplementary Figure S6**
**McKenney et al.**

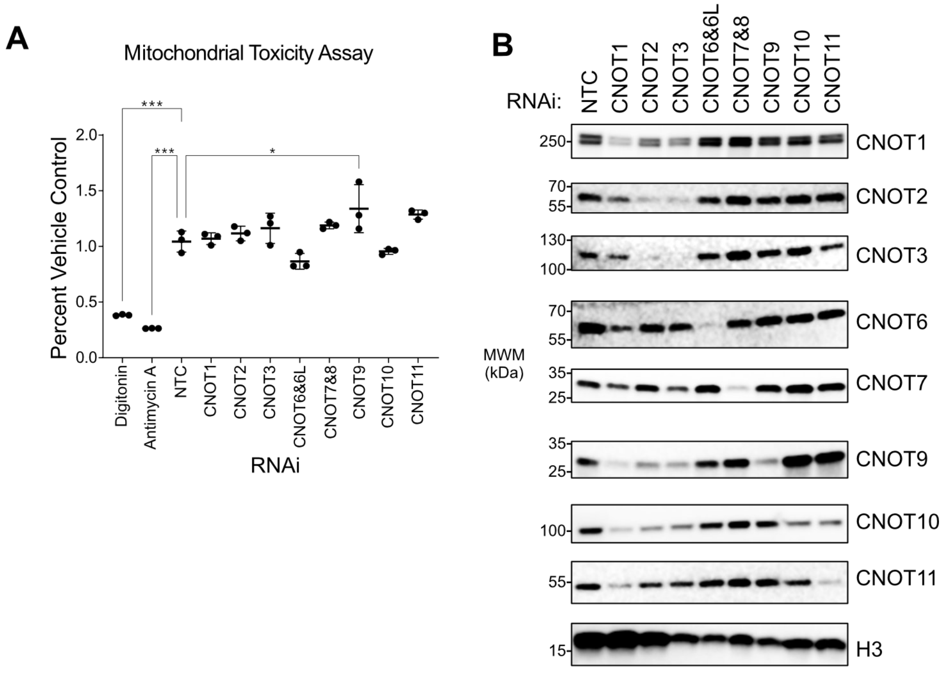

**Figure S6. Assessment of mitochondrial function following CCR4–NOT subunit knockdown.** (**A**) Mitochondrial function in cells grown in galactose was measured using a mitochondrial ToxGlo assay (Promega) after depletion of individual CCR4–NOT subunits compared to an untreated control (percent vehicle control). Digitonin served as a positive control for membrane integrity and antimycin A for mitochondrial toxicity. n=3 biological replicates; +/- standard deviation (SD). For significance calling, $p < 0.05$ = *, $p < 0.01$ = **, $p < 0.001$ = ***, $p < 0.0001$ = **** based on ordinary one-way ANOVA and Tukey test for multiple comparisons. (**B**) Western blot corresponding to samples used in the mitochondrial Toxglo assay shown in panel (A).

### Supplementary Figure S7
### McKenney et al.

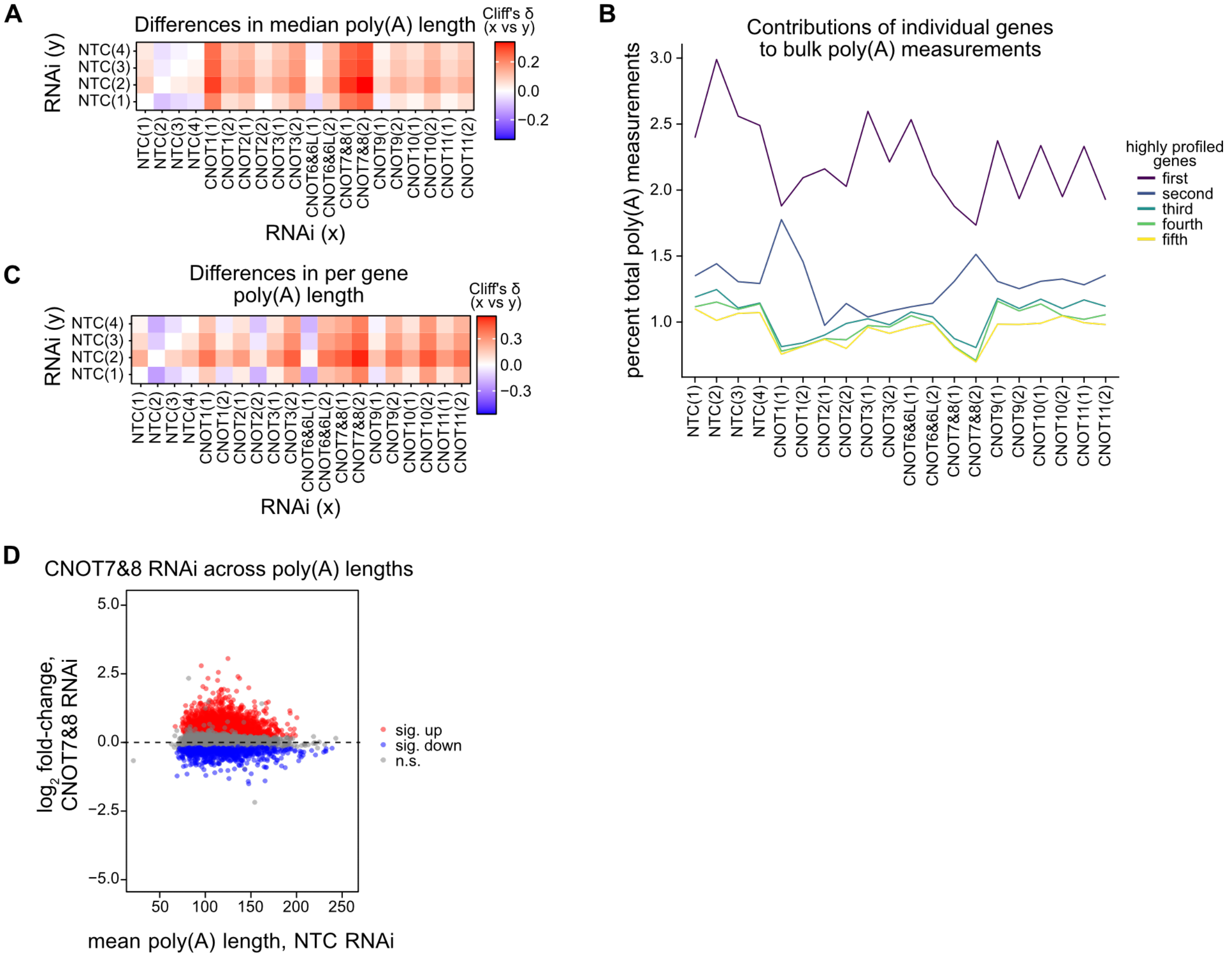

**Figure S7**. **Bulk and gene-level poly(A) measurements**. (**A**) Heatmap of Cliff's delta (δ) between bulk poly(A) measurements. (**B**) Percentage of total poly(A) measurements, per sample, for the five most highly profiled genes. (**C**) Heatmap of Cliff's delta (δ) between mean gene-level poly(A) measurements. (**D**) Comparison of mean poly(A) length in NTC samples and $\log_2$ fold-change in mRNA level following CNOT7&8 depletion.

**Supplementary Figure S8**
**McKenney et al.**

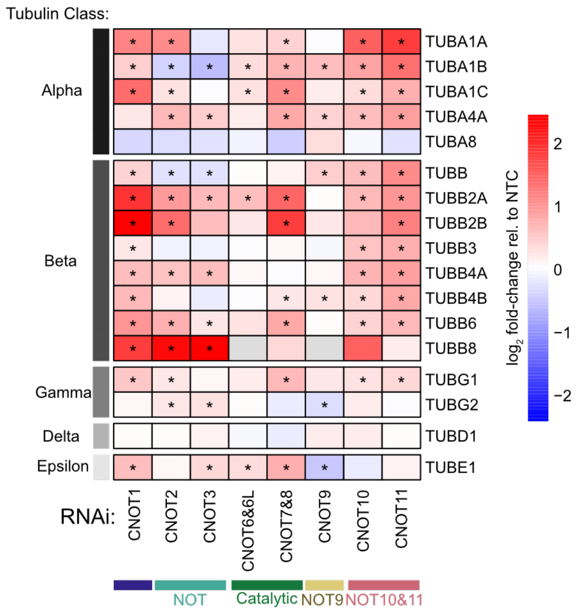

**Fig S8. Tubulin class-specific expression changes following CCR4–NOT depletion.** Heat map showing $\log_2$ fold-changes for 17 tubulin genes detected in HCT116 cells after knockdown of individual CCR4–NOT subunits. Genes are grouped and color-coded according to tubulin class. Asterisks indicate statistically significant changes, defined as $p^{adj} < 0.05$. Note that *TUBA3D*, *TUBA3E*, *TUBA4B*, and *TUBB1* were not expressed in HCT116 cells and thus were not included.

### Supplementary Figure S9
McKenney et al.

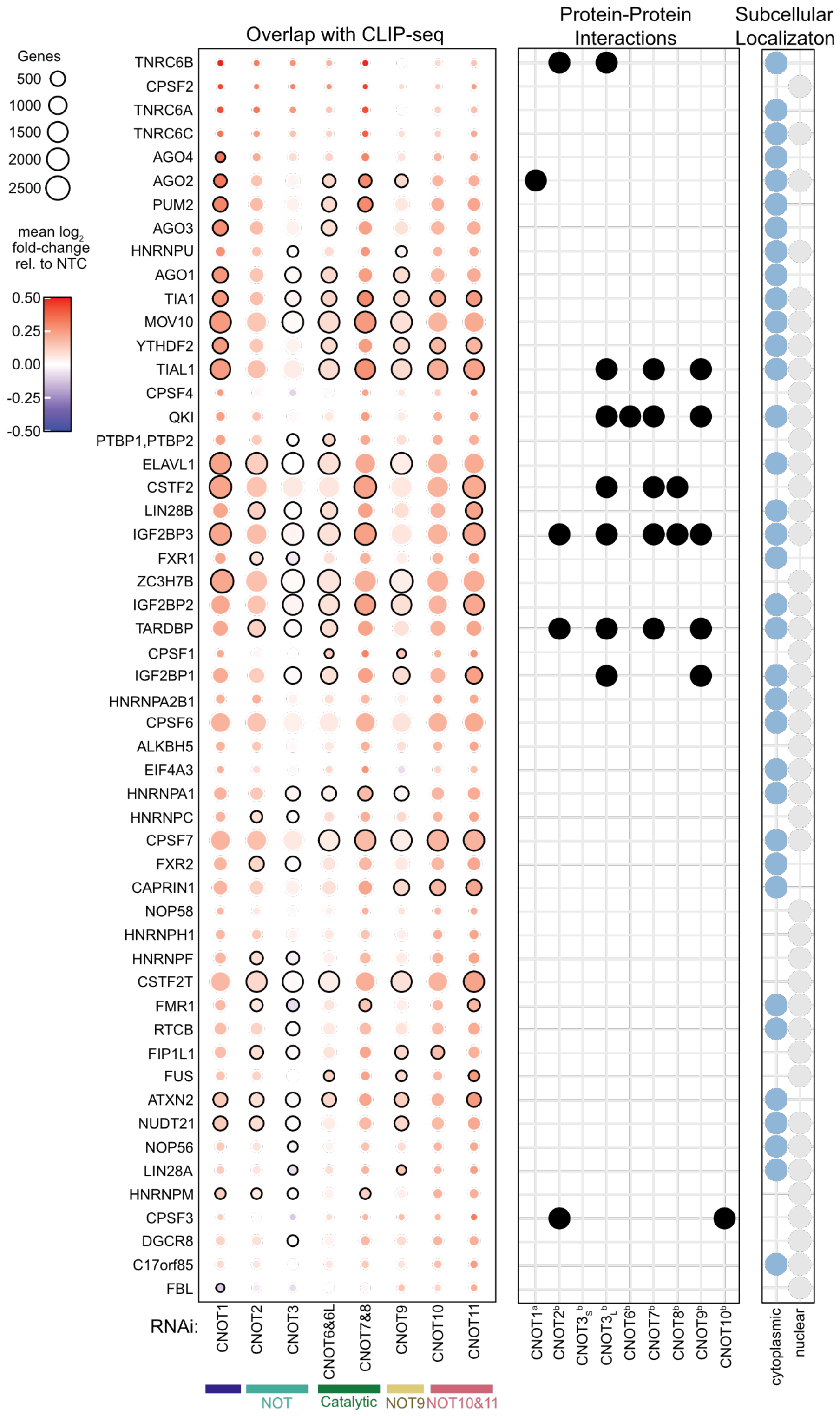

**Fig S9. RNA-binding proteins associated with CCR4–NOT-regulated transcripts.** Dot plot showing the overlap between transcripts upregulated upon knockdown of individual CCR4–NOT subunits and mRNA targets of 53 RNA-binding proteins (RBPs), identified by CLIP-seq in HEK293T cells, as in **Fig. 7A**. Dot size corresponds to the number of overlapping genes and outlined dots indicate statistical significance ($p<0.05$) based on Benjamini–Hochberg-adjusted p-value from a two-sided Wilcoxon test comparing target $\log_2$ fold-change values to all genes not bound by that RBP. Color denotes the mean $\log_2$ fold-change of these genes following knockdown of the relevant subunits, relative to NTC controls. Known protein-protein interactions from (a) Kulkarni (2025) or (b) Lau (2009) are marked with black dots. Subcellular localizations curated from UniProt, cytoplasmic (blue) or nuclear (grey), are indicated.

**Supplementary Figure S10**
**McKenney et al.**

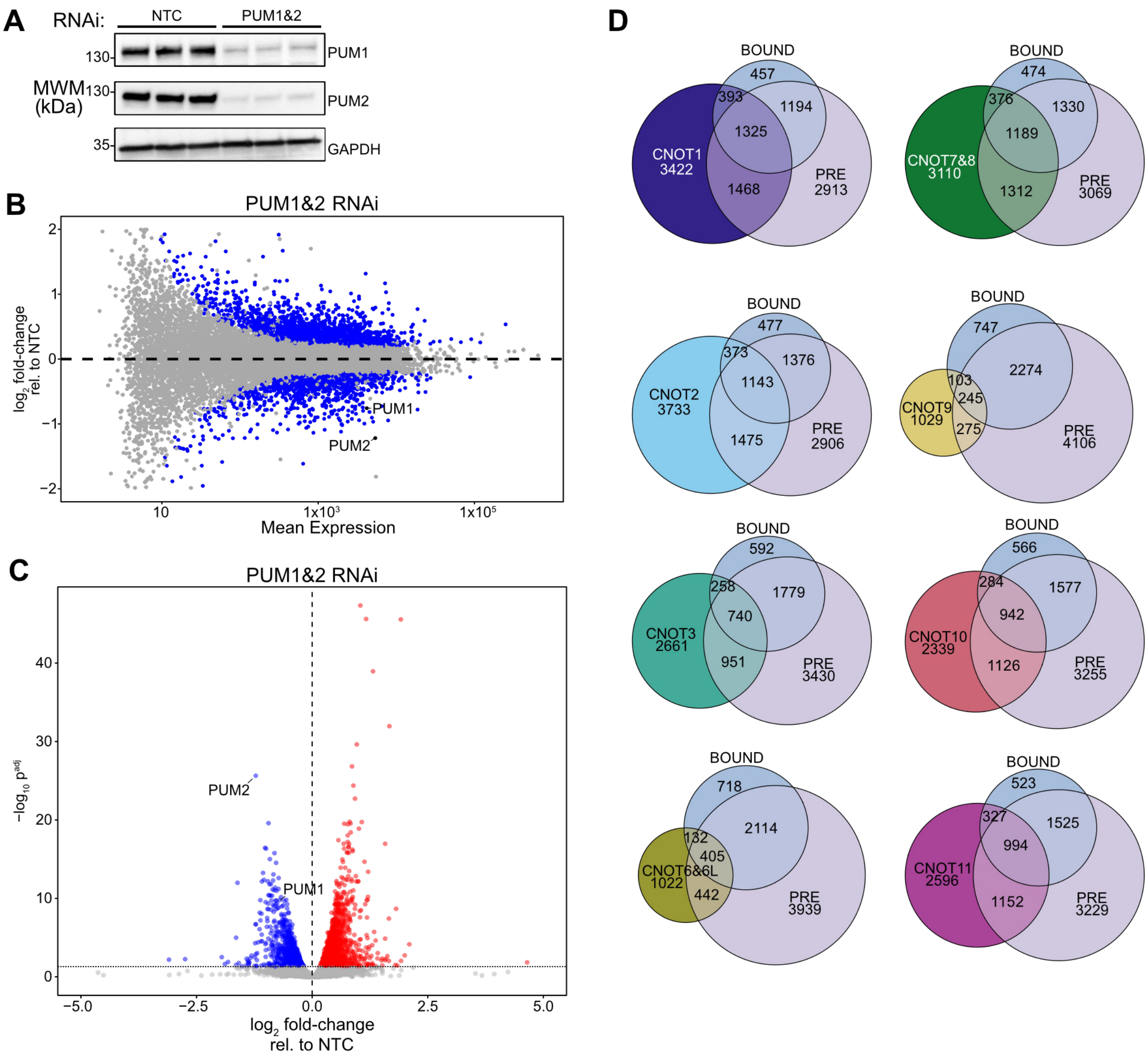

**Fig S10. Effect of CCR4–NOT and Pumilio knockdowns on the transcriptome.** (**A**) Western blot confirming knockdown of PUM1 and PUM2, with GAPDH as a loading control. (**B**) MA plot showing $\log_2$ fold-change versus mean expression for PUM1&2 RNAi relative to the non-targeting control (NTC). Significant changes are shown in blue. (**C**) Volcano plot showing Benjamini–Hochberg-adjusted p-value ($p^{adj}$) versus $\log_2$ fold-change. Significantly downregulated, upregulated, and nonsignificant changes are shown in blue, red, and gray, respectively. PUM1 and PUM2 are indicated. (**D**) Venn Diagrams showing overlap among PRE-containing transcripts, PUM-bound targets, and transcripts significantly upregulated in each CCR4–NOT subunit knockdown.

### Supplementary Figure S11
McKenney et al.

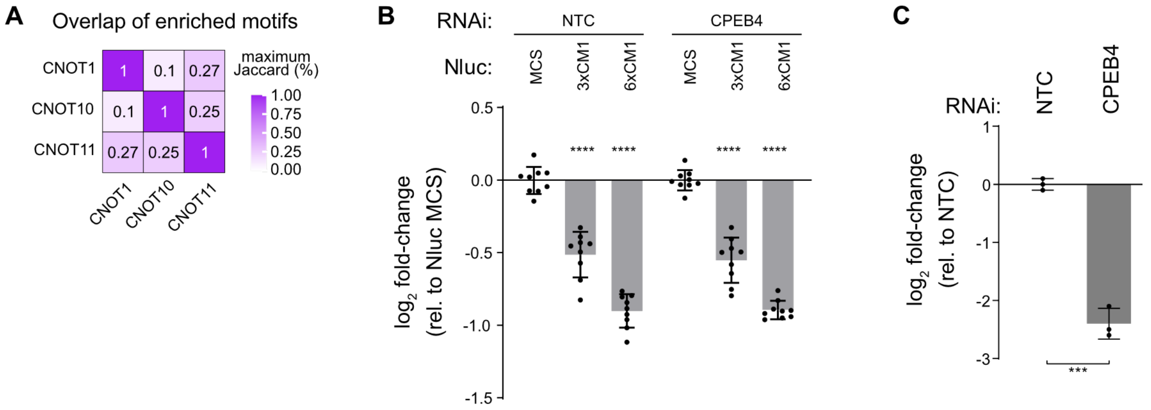

**Fig S11. Similarity and functional impact of CNOT1, CNOT11, and CNOT10 motifs identified by FIRE.** (**A**) Overlap among FIRE-identified motifs for CNOT1, CNOT10, and CNOT11, with numbers indicating percentage of the maximum possible Jaccard value. (**B**) Repressive activity of the CCR4–NOT complex measured in HCT116 cells using Nano-luciferase reporters containing three or six motifs (Nluc 3xCM1 and 6xCM1, respectively) identified by FIRE within a minimal 3′UTR, relative to a control reporter containing a 3′UTR multiple cloning site (MCS). Reporter activity was quantified under non-targeting control (NTC) conditions and following RNAi knockdown of CPEB4. Firefly luciferase (Fluc) served as an internal control to assess variability in transfection efficiency. n=9; 3 experiments, each with 3 biological replicates; +/- standard deviation (SD). Significant differences determined by ordinary one-way ANOVA and Tukey test for multiple comparisons. (**C**) Verification of CPEB4 by RT-qPCR . n=3 biological replicates; +/- SD. Significant differences between NTC and CPEB4 RNAi determined via an unpaired student's t-test. In all panels, $p < 0.05$ = *, $p < 0.01$ = **, $p < 0.001$ = ***, $p < 0.0001$ = ****.
