## Supplementary File S3 for "Global Regulatory Functions of the Human CCR4–NOT Complex in mRNA Deadenylation and Decay"

ENCODE Guidelines and Best Practices for RNA-Seq: Revised December 2016

**Metadata to be supplied with each RNA-seq experiment:**

1. **Donor Information- HCT116 (CCL-247, ATCC) cell line:**
   a. **Sex:** Male
   b. **Age:** 48 years (adult)
   c. **Ethnicity:** No ethnicity information available
   d. **Familial relationships to other donors:** None reported
   e. **External donor links (if applicable):** Not applicable
2. **Sample Information:**
   a. **Material type (tissue, cell line, primary cells, etc.):** Human immortalized colorectal carcinoma cell line HCT116 (CCL-247, ATCC)
   b. **Ontology term:** EFO:0002824 (HCT116 cell line)
   c. **Treatments or genetic modifications:** RNA interference with CNOT subunit siRNAs or non-targeting control RNAs (see materials and methods- ***siRNA transfection***)
   d. **Subcellular fraction or derivation from another sample:** N/A
   e. **Starting material abundance:** Cell densities ranged between approximately 4-7 x 10^6^ cells per sample with viability >90% as determined by Trypan blue staining using a Countess Automated Cell Counter (Thermo Fisher).
   f. **Lot #, catalog #, supplier:**
      i. **Catalog number:** ATCC CCL-247
      ii. **Supplier:** American Type Culture Collection (ATCC)
      iii. **Lot number:** 70051513
   g. **Culture protocol:** Standard ATCC conditions, see materials and methods- ***Cell Culture.***
   h. **Phenotyping/Characterization:** STR profiling confirmation by ATCC. Mycoplasma testing: none detected.
3. **RNA Information:**
   a. **RNA type:** Total RNA
   b. **RNA size fraction:** Protocol for RNA isolation can be found in materials and methods (***RNA purification*)**. RNA isolation from HCT116 cells was performed using the SimplyRNA cells kit and Maxwell RSC instrument (Promega) to purify total RNA without a size selection. According to Promega, the kit is biased to recover RNAs larger than 100 nt.
   c. **rRNA depletion:** RiboZero Plus (Illumina)
4. **Protocol Documentation:**
   a. **RNA isolation method:** Protocol for RNA isolation can be found in materials and methods (RNA purification). RNA isolation from HCT116 cells was performed using the SimplyRNA cells kit and Maxwell RSC instrument (Promega).
   b. **Size selection method:** No size selection was performed.
   c. **rRNA removal method:** This was performed using the Stranded Total RNA Library Prep with RiboZero Plus kit (Illumina) by the University of Minnesota Genomics Center.
   d. **Oligo-dt selection method:** N/A
   e. **DNase treatment method:** DNase treatment was performed during the Maxwell purification where twice the amount of DNase was added for the on-bead DNase treatment.
5. **RNA Quantification and Quality Control:**
   a. **Total RNA QC results:** The concentration of total RNA was determined by UV absorbance using a Nanodrop spectrophotometer (Thermo Fisher Scientific) and the quality of the RNA was determined by Agilent Tapestation RNA ScreenTape analysis yielding RNA integrity number (RIN) values ≥9.9 for all samples.
   b. **Post-oligo-dt selection QC results:** Not applicable
   c. **Post-rRNA depletion QC results:** According to Illumina FastQC sequence content analysis, <0.02% of reads aligned to rRNA reference sequences, confirming efficient rRNA depletion.
   d. **Post-library construction QC results:** All libraries passed quality control. Library concentration using PicoGreen. Fragment size distribution and quality were assessed using Agilent TapeStation D1000 by the University of Minnesota Genomics Center.
6. **Library Construction:** Libraries were prepared using the Stranded Total RNA Library Prep with RiboZero Plus kit (Illumina) by the University of Minnesota Genomics Center**.**
7. **Sequencing Information:**
   a. **Platform:** NovaSeq S4 (Illumina)
   b. **Read format:** Paired-end
   c. **Read length:** 150 bp
   d. **Barcode placement and sequence:** Standard (Illumina)
   e. **Custom primer sequences:** Not applicable

**RNA Sequence Experiment Design: Replication, sequencing depth, spike-ins**

1. **Replication:**
   a. **Number of biological replicates:** Three biological replicates per knockdown condition.
   b. **Number of technical replicates:** Not applicable
   c. **Replicate concordance (Spearman correlation):** Spearman correlation coefficients >0.98 across all biological replicates.
2. **Sequencing Depth:**
   a. **Aligned reads per library:** Each library yielded an average of >76 million uniquely aligned read-pairs per sample.
3. **Spike-ins:** External RNA Controls Consortium (ERCC) Spike-In Control Mixes were included for normalization in all samples at 2 ul of a 1:1000 dilution for 100 ng total RNA input before library preparation according to manufacturer's protocol (Thermo Fisher Scientific, 4456740).

**Post-Sequencing Quality Control**

1. **Mapping and Alignment:**
   a. **Reference genome used:** UCSC hg38 genome + ERCC92 spike-in sequences
   b. **Annotation set included:** hg38 UCSC refGene + ERCC92 spike-in annotations
   c. **Spike-in sequences included in reference:** Yes
   d. **Mapping software and version:** Nextflow RNA‑seq pipeline (v3.9) using STAR
